# Oxysterol accumulation in aging cells alters GPCR signalling

**DOI:** 10.1101/2024.05.09.593443

**Authors:** Suramya Asthana, Anant Verma, Baivabi Bhattacharya, Arnab Nath, Nithin Sajeev, Kiran Maan, Raji R. Nair, K. Ganapathy Ayappa, Deepak K. Saini

## Abstract

Organismal aging is accompanied by the accumulation of senescent cells in the body, which drives tissue dysfunction. Senescent cells have a distinctive profile, including proliferation arrest, resistance to apoptosis, altered gene expression, and high inflammation. Despite global signalling and metabolic dysregulation during senescence, the underlying reasons for changes in signalling remain unclear. GPCRs are pivotal in cellular signalling, dynamically mediating the complex interplay between cells and their surrounding environment to maintain cellular homeostasis. The chemokine receptor CXCR4 plays a crucial role in modulating immune responses and inflammation. It has been shown that expression of CXCR4 increases in cells undergoing senescence, which enhances inflammation post-activation. Here we examine CXCR4 signalling in deeply senescent cells, where cholesterol and its oxidized derivatives, oxysterols, affect receptor function. We report elevated oxysterol levels in senescent cells, which altered classical CXCL12-mediated CXCR4 signalling. Tail-oxidized sterols disrupted signalling more than ring-oxidized counterparts. Molecular dynamics simulations revealed that 27-hydroxycholesterol displaces cholesterol and binds strongly to alter the conformation of critical signalling residues to modify the sterol-CXCR4 interaction landscape. Our study provides a molecular view of the observed mitigated GPCR signalling in the presence of oxysterols, which switched G-protein signalling from Gα_i/o_ to Gα_s_ class. Overall, we present an altered paradigm of GPCR signalling where cholesterol oxidation alters the signalling outcome in aged cells.

**Significance Statement:** Our study brings to light a novel and significant discovery in aged cells: the accumulation of oxysterols, oxidized forms of cholesterol, critically impairs CXCR4-dependent signalling and alters G-protein coupling specificity. This effect of oxysterols is demonstrated for the first time in aged cellular models, providing a molecular basis for a multitude of observed alterations in senescence, such as compromised immune functions and a decline in cellular responsiveness with age. Our research not only fills a crucial gap in understanding the aging process at the molecular level but also identifies potential targets for therapeutic interventions aimed at mitigating age-related cellular dysfunctions and diseases.

## Introduction

Aging is defined as the temporal deterioration of physiological functions of organisms. Despite significant progress in extending the human lifespan through modern medicine, improved sanitation, and nutrition, the undesirable effects associated with aging have not been alleviated[1]. There is increasing evidence that aging occurs at the cellular level and cellular senescence is a significant contributing factor. It is now well established that accumulation of senescent cells over the years results in tissue dysfunction leading to organismal aging[2]. Thus, it is crucial to understand the molecular basis of aging and identify possible therapeutic intervention approaches and targets.

GPCR super-family of transmembrane proteins are involved in regulating various cellular processes in response to a large variety of extracellular cues, which include light, hormones, amines, etc., and are widely used as drug targets in almost all human disorders except aging, where no unique druggable GPCR is reported till now. CXCR4, a chemokine GPCR, is ubiquitously expressed, and CXCL12 or SDF-1α is the only known ligand for this receptor. This receptor is reported to be upregulated in aging[3, 4] and senescent cells[5, 6]. The canonical G-protein-dependent pathway mediated by CXCR4 inhibits adenylyl cyclase through the Gα_i/o_ subunit, decreasing cellular cyclic AMP (cAMP) levels. It activates phospholipase Cβ (PLCβ) and phosphoinositide-3 kinase (PI3K) through Gβγ subunits triggering calcium release from ER stores as well as drives non-canonical MAPK activation[7, 8]. Interestingly, in a genetic disorder, WHIM syndrome, where a C-terminal truncation mutation that constitutively activates the receptor is reported, signs of accelerated aging are observed[9]. In addition, several types of cancer cells express the CXCR4 receptor, and expression levels are negatively correlated with survival. CXCR4 expression promotes proliferation, survival, and metastasis of cancer stem cells[5, 10].

In GPCRs, signal transduction occurs primarily through the transmembrane helices involving both the extracellular and intracellular loops that bind ligand and mediate G-protein coupling. Given this, signalling is sensitive to changes in the lipid environment, dominated by cholesterol-protein interactions, as shown for a wide variety of GPCRs[11–13]. CXCR4 has been shown to co-localize in lipid rafts associated with membrane domains rich in cholesterol and sphingomyelin[14]. Although elevated levels of oxysterols, a naturally occurring form of cholesterol, have been implicated in pathologies such as cardiovascular diseases, autoimmune disorders and various metabolic disorders, its influence on GPCR, CXCR4-dependent signalling, the focus of this manuscript, is poorly understood[15].

Membrane oxysterols modulate lipid packing, influence microdomain formation[16] and can potentially alter the protein-sterol binding landscape. Molecular dynamics (MD) simulations have been widely used to study several aspects of GPCR activation and dynamics on interaction with cholesterol[17–19]. Using a combination of experiments and all-atom MD simulations, we explore the altered protein-sterol interaction landscape to explain the influence of oxysterols on CXCL12 mediated CXCR4 signalling.

We have assessed the signalling after the external addition of oxysterols or depletion of cholesterol as well as during senescence. For external addition experiments, two ring oxidized, 7β-hydroxycholesterol and 7-ketocholesterol and four tail oxidized cholesterols, 25-hydroxycholesterol, 27-hydroxycholesterol, 22(R)-hydroxycholesterol, and 24(S)-hydroxycholesterol were used, which disrupted GPCR signalling. To support experimental findings, fully atomistic 3 µs long MD simulations of CXCR4 in POPC:Cholesterol (80:20) and membranes with 10% oxysterols were used to differentiate residue-wise sterol binding lifetimes for the 7 transmembrane helices in the presence of oxysterols. Along with the conformational changes and sterol binding propensities at the critical signalling residues, the MD simulations show that the most significant perturbation in sterol binding patterns occurs for the tail oxidized sterols supporting the disrupted calcium signalling patterns observed in the experiments. This is the first study of its kind where the molecular aspects of oxysterol-protein interactions that influence GPCR signalling pathways during cellular senescence are analyzed. This study also provides a framework for the observation of altered therapeutic outcomes for GPCR targeting drugs in aged cells.

## Results

### CXCR4 signalling is altered during senescence

The CXCR4 receptor has been reported to be upregulated in various models of cellular senescence and naturally aged neutrophils[20]. In this study, the genotoxic stress-induced model of senescence was used, where the HeLa cells were treated with 5-bromo-2’-deoxyuridine (BrdU) or ionizing radiation and incubated for 4 days, followed by analysis to ensure senescence induction. The senescence state was confirmed by assessing several markers, including the presence of a large, flattened morphology in vitro and positive stain for SA-βgal (Figure 1a); an increase in the number of γH2AX foci per cell in treated cells, confirming the presence of DNA damage (Figure 1b). The expression of p21 and CXCR4 was also high in treated cells, confirming the establishment of senescence (Figure 1c, d). Previously shown upregulation of the CXCR4 receptor was also validated by surface staining for the CXCR4 receptor (Figure 1e). The signalling through this receptor is mediated by the ligand CXCL12 and is involved in diverse functions like cell proliferation and migration and enhancing inflammation. Given that the ligand is not expressed by HeLa cells, it makes it an appropriate model system to study signalling related changes by external ligand addition.

**Figure 1:**
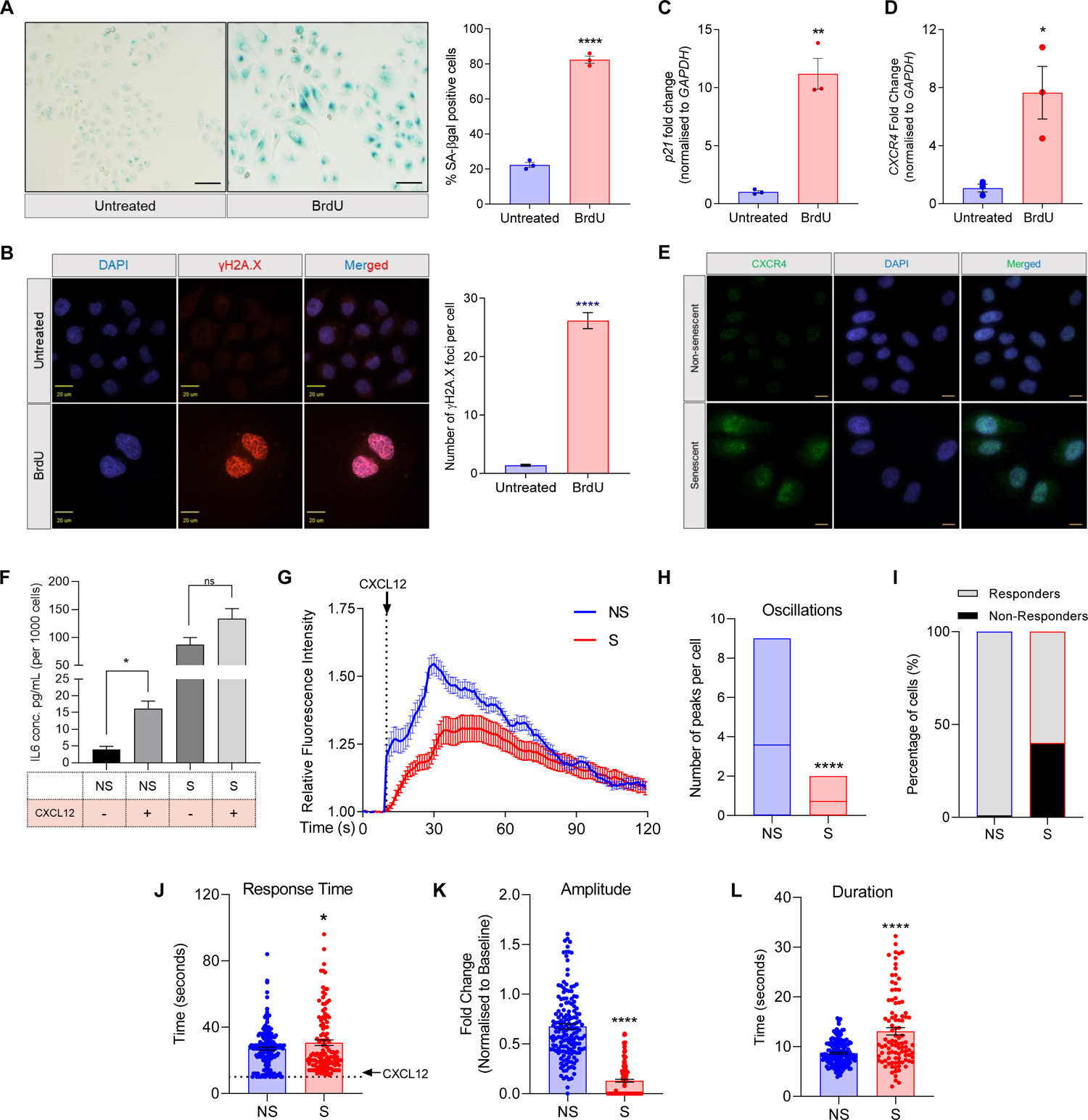
Cellular senescence and CXCR4 signalling. HeLa cells were treated with BrdU for 96 hours to induce senescence. (a) SA-βgal staining, left - untreated control, right - BrdU treated cells. The graph reports quantification of SA-βgal positive cells. (b) Immunofluorescence analysis of γH2AX foci formation. The left - representative images, and right – quantification of number of foci per cell. Gene expression quantitation of (c) p21 and (d) CXCR4. (e) Surface immunostaining of CXCR4 receptor. (f) Quantification of secreted IL6 levels in various conditions as listed. Non-senescent (NS) and senescent (S) cells. (g) Calcium release in response to CXCR4 activation (Movie S1, S2). Single-cell analysis of calcium response, (h) mean oscillations, (i) the number of responding cells, (j) response time; (k) amplitude of the response; and (l) response duration (n=3; N>100 for all groups)

We tested the various signalling outcomes in the deeply senescent cells and found that the inflammatory response was impaired upon stimulation (Figure 1f), unlike what is recorded during the onset of senescence. Further investigation revealed that the calcium response mediated by the CXCL12-CXCR4 pathway activation was also impaired, and the mean number of calcium peaks was lower in senescent cells (Figure 1g, h). Single-cell analysis revealed that a significant percentage of the population did not elicit any response, and the occurrence of calcium peaks was also delayed (Figure 1i, j; S1). In responding senescent cells, the response amplitude was much lower (Figure 1k), and the average duration of sustenance for each peak was higher (Figure 1l). The signalling readouts were found to be similar in the ionizing radiation model of senescence (Figure S1). These findings indicated that the CXCR4 signalling was impaired in deeply senescent cells.

### Gene expression analysis reveals alteration in CXCL12-CXCR4 signalling in the senescent state

To decipher the mechanism behind the altered signalling axis, we performed comparative gene expression analysis of non-senescent and senescent cells, in the basal and CXCL12 stimulated conditions. The response index (RI) to CXCL12 in non-senescent (RINS) and senescent (RIS) cells was calculated by normalizing the stimulated to the basal state (Figure 2a). Based on the RI, the differentially expressed genes were segregated into four categories, as shown in Figure 2b. Gene set enrichment analysis (GSEA) revealed pathways that were affected in senescent cells compared to non-senescent cells (Figure 2c). While immune system processes, cytokine mediated signalling and GPCR signalling were among the top altered pathways, the appearance of phospholipid biosynthetic processes in this list raised an interesting question about altered lipid content being responsible for altered signalling. We also performed GSEA for non-senescent and senescent samples in the basal state, and among the top pathways, cholesterol homeostasis and organic hydroxy compound transport pathways were upregulated (Figure 2d). Cholesterol is an essential part of the plasma membrane and crucial for signalling events; it has been extensively associated with the activity of many receptors[11, 17, 21, 22] including CXCR4[14, 23]. To investigate the role of cholesterol, if any, we first measured the intracellular cholesterol levels by LC-MS/MS and found no significant differences in the total cholesterol levels (Figure 2e). However, feedback from oxysterols (oxidized cholesterol) regulates cellular cholesterol homeostasis, which involves cholesterol efflux, esterification, bile acid synthesis etc[24]. Oxysterols are physiologically generated by enzymatic or non-enzymatic pathways, including oxidation by reactive oxygen species (ROS)[25]. It is well established that senescent cells have high ROS, that we observed in our system as well (Figure 2f); concomitant to this, we also found enriched ROS metabolism pathways in GSEA (Figure 2d). This consequently raised an intriguing possibility about the direct impact of potentially elevated levels of oxysterols in altered GPCR signalling in aged cells.

**Figure 2:**
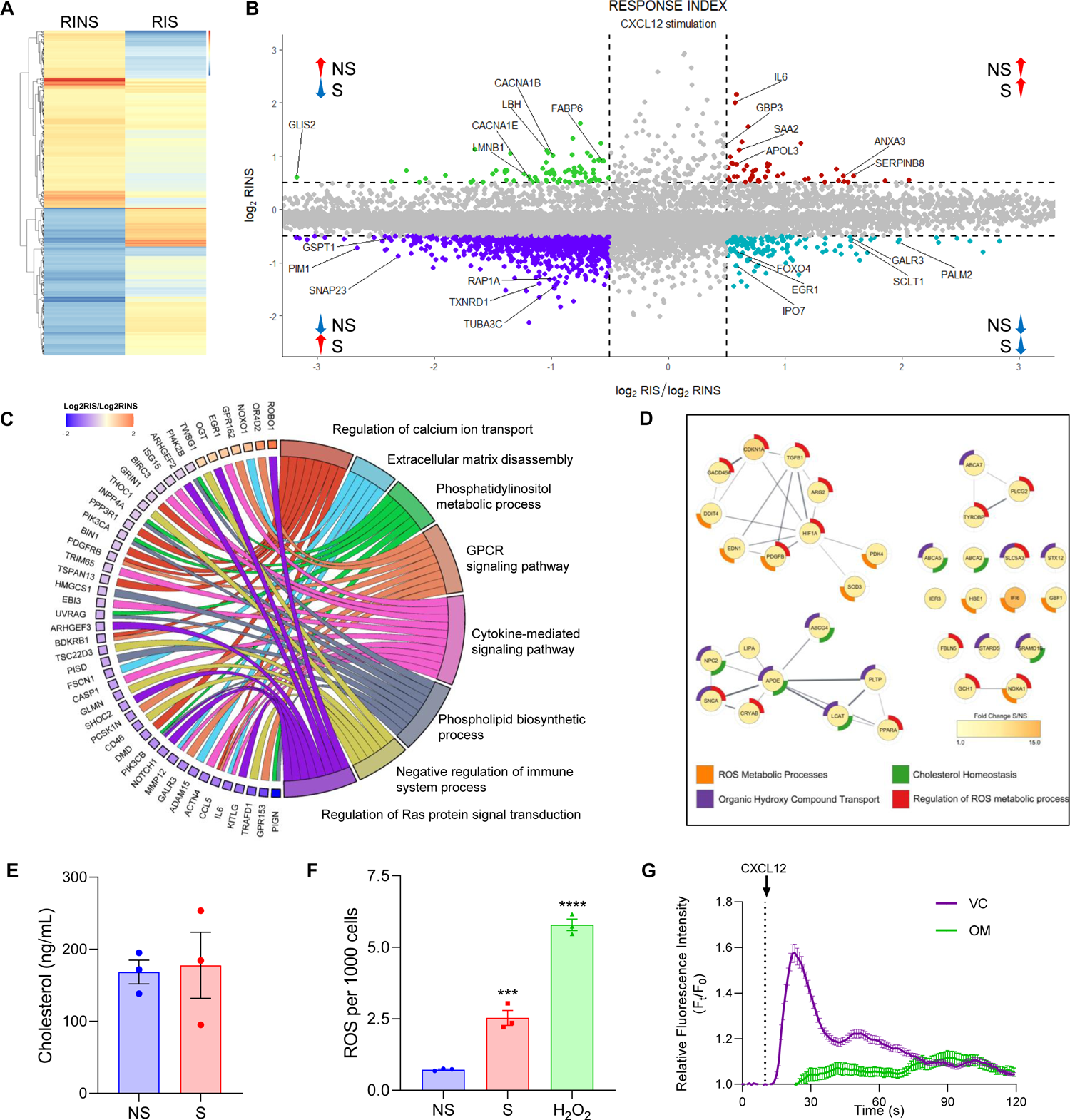
Identification of physiological changes during senescence. NS and S HeLa cells were stimulated with the ligand CXCL12, and microarray analysis was performed. (a) Heat map of Response Index (RI) in non-senescent (RINS) and senescent (RIS) cells (n=2). (b) Genes differentially regulated in presence of CXCL12 clustered into four categories (represented in each quadrant). (c) Gene set enrichment analysis (GSEA) for top pathways after stimulation in senescent and non-senescent cells. (d) Network map for selected top pathways in GSEA of non-senescent and senescent cells at the basal state. (e) Intracellular cholesterol content measured by LC-MS/MS analysis. (f) Intracellular ROS levels measured by DCFDA. H2O2 treatment is used as a positive control. (g) Calcium response in non-senescent cells treated with vehicle (VC) or oxysterol mixture (OM) after stimulation with CXCL12. (n=3; N>100 for all groups)

To first test if oxysterols can affect GPCR signalling, non-senescent cells were treated with a mixture of oxysterols containing 2 types of ring (7-BHC and 7-KC) and 4 types of tail (25-OHC, 27-OHC, 22(R)-OHC and 24(S)-OHC) oxidized cholesterols (Figure S3) and the calcium response after CXCR4 activation was recorded. Oxysterol treated non-senescent cells also showed an impaired calcium response (Figure 2g) and the loss of oscillations and delayed response was similar to the patterns seen in senescent cells.

### Altered membrane composition affects CXCR4 signalling during senescence

Based on these observations, we decided to measure if the oxysterol levels in non-senescent and senescent cells differ. We found that the levels of both ring and tail oxidized oxysterols were significantly higher in senescent cells (Figure 3a; S4a). To further validate this, we measured the levels of 27-hydroxycholesterol (27-OHC) in senescent primary immortalized cell lines of different origins, which were found to be significantly higher (Figure S4b-c), indicating that the presence of oxysterols is a senescence-specific alteration, independent of the cell type and mode of senescence induction.

**Figure 3:**
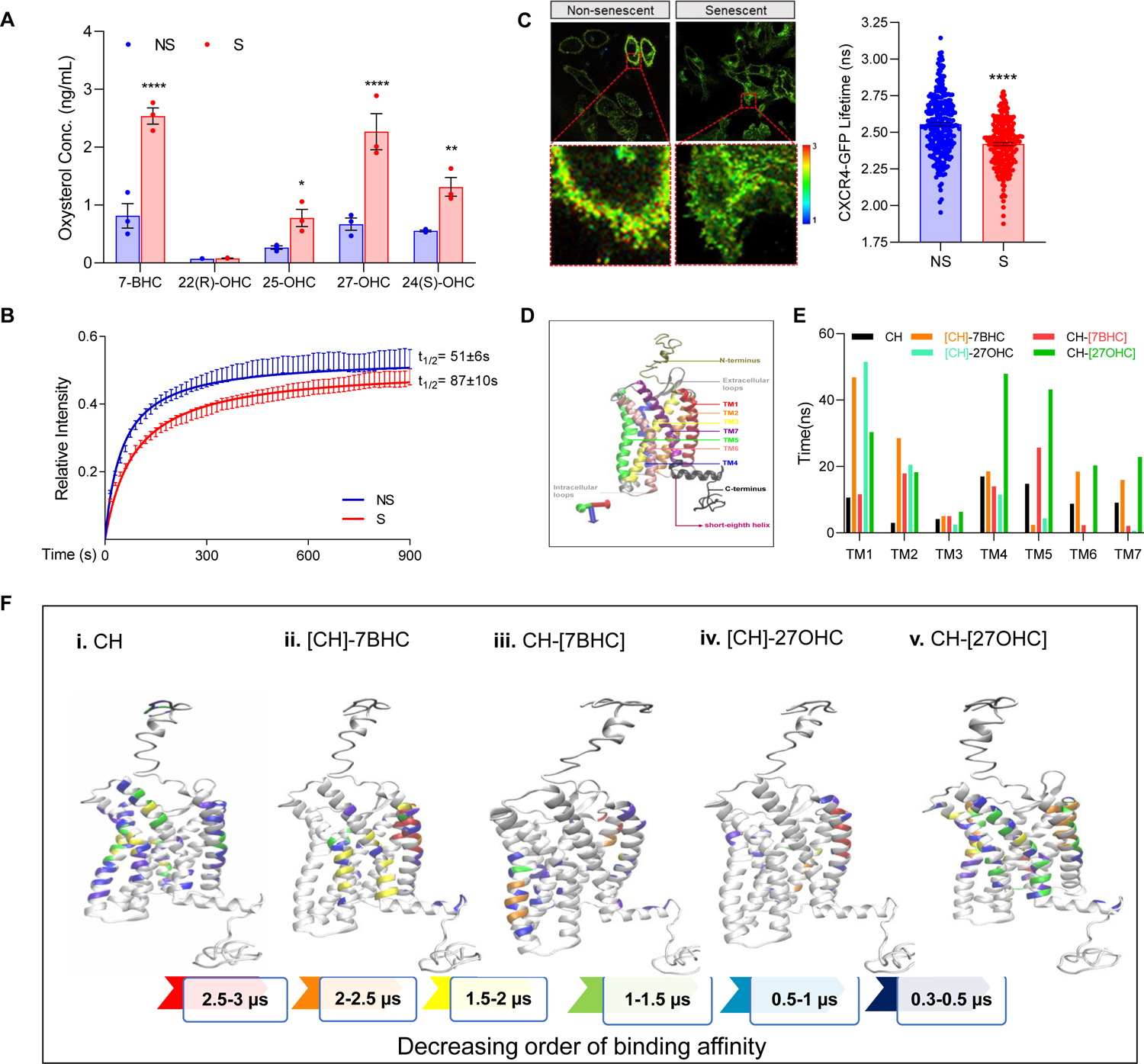
Effect of oxysterol accumulation in senescent cells on CXCR4 receptor. (a) Measurement of oxysterol levels by LC-MS/MS analysis. (b) CXCR-GFP FRAP curves in non-senescent cells (n=6; N=91), senescent cells (n=4; N=99); t1/2 is denoted (n=3; N>100 for all groups). (c) FLIM analysis of CXCR4-GFP. Left - representative images, and right – distribution of lifetime values. (d) Structure of CXCR4 used in simulations. Different domains are highlighted in different colours. (e) TM-wise binding of sterols calculated as the cumulative binding time per sterol molecule, divided by the number of residues in a TM. (f) Binding affinity of [CH] (j-i), [CH]-7BHC(j-ii), CH-[7BHC] (j-iii), [CH]-27OHC (j-iv) and CH-[27OHC] (j-v) with CXCR4. Decreasing affinity of binding is represented by color index shown. [] refers to the component in the binary mixture depicted in the figure.

It is well established that the CXCR4 receptor on the cell membrane is enriched in lipid rafts and undergoes oligomerization upon ligand stimulation[26]. We thus evaluated the effect of altered membrane dynamics on CXCR4 receptor mobility and oligomerization on senescent membranes. Towards this, we performed FRAP using CXCR4-GFP, which showed lower recovery time in senescent cells, indicating reduced receptor mobility on the membrane in senescence (Figure 3b). Receptor oligomerization status was analyzed using fluorescence lifetime imaging with CXCR4-GFP, and it was found that the lifetime was significantly lower in senescent cells, as well as non-senescent cells treated with oxysterol mixtures compared to controls (Figure 3c; S5), indicative of reduced receptor clustering. Given that the signalling was impaired, we checked the activation status of the receptor by monitoring its internalization post stimulation. The receptor internalization was not affected in senescent cells (Figure S6a) nor oxysterol treated non-senescent cells (Figure S6b) as observed from the appearance of internalized receptor punctae after stimulation. The observed changes in CXCR4 signalling due to the presence of oxysterols in the membrane thus led us to question the specific role of oxysterols and their impact on the CXCR4-sterol interactions in the membrane.

### Molecular Dynamics simulations for cholesterol-oxysterol interaction with CXCR4 in the membrane

To understand the impact of enhanced oxysterols on receptor structure and associated function, all-atom MD simulations of CXCR4 (Figure 3d) in a phospholipid bilayer with different oxysterol compositions (Table 1) were carried out and comparisons made with membranes containing only cholesterol. The binding times for all 352 residues in the receptor over the entire 3 μs simulation for the different cholesterol systems were recorded (Figure S7). The most significant modification was observed with 27-OHC binding sites that span the entire protein surface. It displaced cholesterol in helices TM4-TM7 with a near complete replacement in helices TM6 and TM7 (Figure 3e), implicated in downstream signalling[27]. Residue-wise binding hot spots of oxysterols (Figure 3f) in CXCR4 show that cholesterol binding in the CH membrane is much weaker than oxysterol binding in CH-27OHC membranes. In contrast to 27-OHC, the binding of 7-BHC was considerably reduced, and an increase in CH binding was observed in the CH-7BHC membranes. Interestingly, 27-OHC samples the membrane in parallel and perpendicular orientations over the course of the 3 *µ*s simulation (Figure S8). This is in sharp contrast to cholesterol or 7-BHC, where predominantly membrane perpendicular orientations were sampled. The simulations provide a molecular view of the altered cholesterol-CXCR4 binding landscape in the presence of oxysterols for the first time. Combined with the experimental observations, our findings suggest that the presence of oxysterols in senescent membranes could potentially alter CXCR4 signalling.

**Table 1:** Details of MD simulations with CXCR4 indicating system name, the membrane, and run time of all three systems, along with the sterol abbreviations used in the manuscript. Square brackets [] are used to denote the specific sterol component in the mixture.

| System Name | Membrane Composition | Run Time | Sterol abbreviations |
| --- | --- | --- | --- |
| CH | 80% POPC, 20% cholesterol | 3 μs | chol ≡ [CH] |
| CH-7BHC | 80% POPC, 10% 7-BHC,<br>10% CHOL | 3 μs | chol ≡ [CH]-7BHC<br>oxysterol ≡ CH-[7BHC] |
| CH-27OHC | 80% POPC, 10% 27-OHC,<br>10% CHOL | 3 μs | chol ≡ [CH]-27OHC<br>oxysterol ≡ CH-[27OHC] |

### Both cholesterol depletion and oxysterols insertion disrupt CXCR4 signalling

Membrane cholesterol content plays a critical role in regulating CXCR4 signalling through direct interactions with the receptor[14, 27]. With this premise, we next examined the effect of cholesterol depletion on CXCR4 signalling by treating non-senescent cells with various concentrations of methyl-β-cyclodextrin to deplete cholesterol, which was confirmed by Filipin staining (Figure S9). On these cells, calcium release was monitored, and stimulation with CXCL12 did not generate an oscillatory calcium response, which was observed in non-depleted cells. Single-cell analysis of calcium oscillations revealed that on average, non-depleted cells showed 4-5 oscillations after stimulation within 2 minutes (Figure 4a, S10a). The calcium response and number of oscillations were impaired after mβCD treatment in a dose-dependent manner, and the number of cells that did not respond to stimulation also increased with cholesterol depletion (Figure 4b). The calcium response was delayed upon cholesterol depletion with 10 mM mβCD, and the response amplitude was lower compared to untreated cells. The response duration, calculated from the full width at half maxima of each peak, was higher in cholesterol-depleted cells, indicating a slower decay in calcium response (Figure S10b). Together, these findings suggest that cholesterol depletion leads to an impaired signalling response.

**Figure 4.**
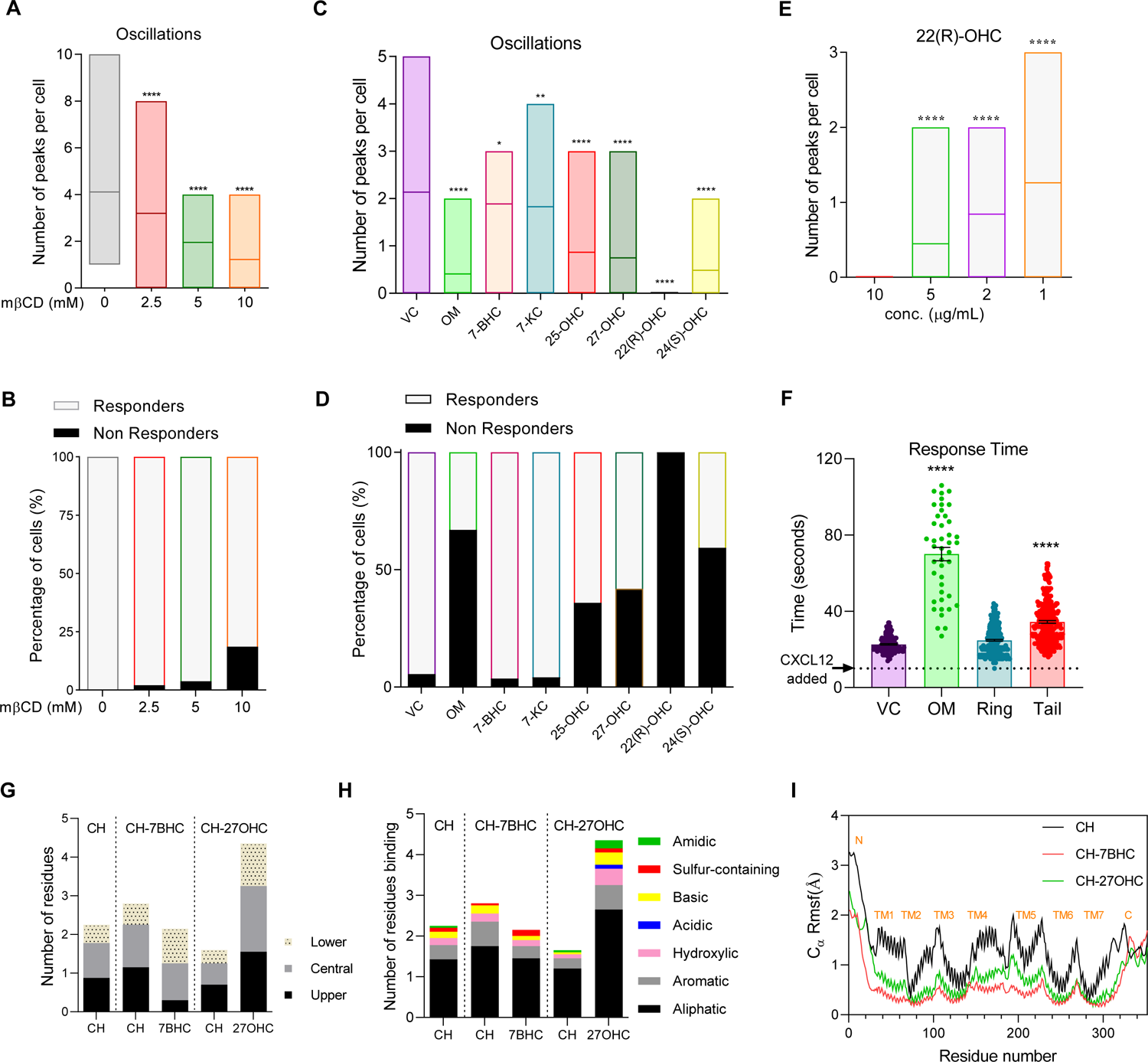
Effect of cholesterol depletion and oxysterol addition on CXCR4 signalling. Single cell analysis of calcium oscillations. (a) Mean number of oscillations after cholesterol depletion (Movie S4). (b) Percentage of cells that responded after cholesterol depletion. (c) Mean number of oscillations in oxysterol-treated cells (Movie S5). (d) Percentage of cells that showed a response in oxysterol-treated cells. (e) Mean number of oscillations in cells after dose titration of 22(R)-OHC. (f) A comparison of response time in tail vs ring oxidized sterol treated cells. (g) Sterol binding to CXCR4 across the z-axis, reported as the number of residues interacting with sterol with a cumulative interaction time greater than 400 ns, divided by number of sterol molecules. (h) Sterol binding to various residues based on their chemical nature in indicated cholesterol-oxysterol systems (indicated on top). Sterol occupation was evaluated as the number of residues binding to the sterol divided by the number of sterol molecules. (i) Root mean square fluctuations (RMSF) of the Cα atoms of CXCR4 in the three cholesterol-oxysterol systems for 3μs simulations.

We next probed the impact of oxysterols on the receptor activity and found that calcium oscillations were lost upon treatment with the oxysterol mixture, as indicated in the previous section. To test the impact of cholesterol oxidation site, cells were treated with individual oxysterols and CXCR4 signalling was evaluated. The ring oxysterols (7-BHC and 7-KC) exhibited a mild reduction in the calcium response, where the majority of cells exhibited a lower number of calcium oscillations. Unlike this, the tail oxysterols (25-OHC, 27-OHC and 24(S)-OHC) drastically affected the calcium response and the mean number of oscillations was significantly lower. No calcium response was observed in 22(R)-OHC treated cells (Figure 4c, S11). Consistent with these observations, the percentage of cells that did not respond to the stimulation was much higher in tail oxysterol treatment, unlike ring oxysterols (Figure 4d). A dose-response analysis of 22(R)-OHC treatment revealed a direct correlation with signalling inhibition (Figure 4e). Single-cell calcium response analysis done only for the cells that showed calcium release in each treatment group showed that the presence of tail oxysterols also delayed the response post-stimulation (Figure 4f). Together, this analysis revealed that oxysterols alter and impair CXCR4-mediated signalling, and more significantly, tail oxysterols have a more substantial impact on the calcium response.

### MD analysis reveals enhanced binding of tail oxysterol with CXCR4 transmembrane helices

Given that the strong effect of tail oxysterol was recorded on CXCR4 activity and the presence of oxysterols modified the binding patterns with the different helices (Figure 3e, f), oxysterol binding patterns were profiled along the membrane. The preferential binding with 27-OHC occurred in protein residues in the upper, central, and lower regions of the membrane (Figure 4g). Furthermore, 39% of the residues binding to 27-OHC were in the central regions of the CXCR4 protein. These effects are partly driven by stronger binding of tail oxysterols with the protein due to the presence of an extra polar group and the tendency of 27-OHC and cholesterol to demix (Figure S12). 27-OHC was found to have a strong binding affinity with hydroxylic, acidic and amidic residues compared to other sterols (Figure 4h). The partitioning of different residue types appeared similar for both cholesterol in CH and 27-OHC in CH-27OHC, indicating competition for similar binding sites. A five-fold increase in H-bonding compared to cholesterol in CH was observed in the cumulative H-bonding time of 27-OHC in CH-27OHC (Figure S12f) and the additional OH in 27-OHC accounts for more than 80% of its H-bonding time with CXCR4 (snapshots illustrating O27 based H-bonds shown in Figure S13). These altered sterol binding patterns appear to directly influence the inherent flexibility of the CXCR4 and oxysterols, resulting in lowered root mean square fluctuations (RMSFs) of the CXCR4 C*_α_* atoms in CH-27OHC and CH-7BHC membranes when compared with the CH only membrane (Figure 4i). Based on these observations, we next examined the influence of oxysterol binding and conformational changes induced in critical signalling residues of CXCR4, which can explain the changes in the output we recorded.

### Oxysterol binding alters the orientation of critical signalling residues

To understand the interaction with critical signalling residues[28], we defined two necessary criteria to decide whether sterol interactions are significantly altered in the CH-7BHC/27OHC systems compared to the CH only system. For the first criterion, sterol molecules must have a cumulative binding time of at least 400 ns with a specific residue and/or its immediate neighbour/s. This was a stringent requirement considering that the cholesterol binding duration observed in many transmembrane proteins was in the order of 100s of ns[21, 29]. The second criterion is related to the altered binding propensities as well as the binding times further sub-classified according to the following conditions for the CH-7BHC/27OHC systems - ( i) enhanced cholesterol-binding (C1): at least a 100% increase in cholesterol binding time compared to CH; (ii) reduced cholesterol-binding (C2): at least 50% reduction in cholesterol binding time compared to CH; (iii) cholesterol replaced with oxysterol (C3): oxysterol binding time is greater than 50 % of the cholesterol-binding time in CH and lower than twice the cholesterol-binding time in CH and (iv) enhanced oxysterol binding (C4): oxysterol binding time increase is greater than twice the binding time of cholesterol in the CH membrane.

The conformational changes of critical signalling residues (Figure S14a) in the presence of oxysterols were quantified using the angle *θ*, between the vector passing through the C-*α* atoms of the first and last residue of the TM helix and the vector passing through the C-*α* atom of the concerned residue and the last carbon atom of the residue (Figure S14b-d). We observed direct enhanced cholesterol binding (C1) where cholesterol remains bound to Y45 for the entire 3 *µ*s in the CH-7BHC system (Figure 5a). In contrast, Y45 is not a binding site for cholesterol in the CH system, and a weak shift of about 5° was observed in *θ* (Figure 5b). An example of reduced cholesterol interactions (C2) was observed at K282 (Figure 5c,d), where a nearly complete cholesterol elimination occurs in CH-7BHC, and 7 - BHC binding does not even occur at this residue. The influence of 27-OHC was more drastic, and a C3-type interaction where 27-OHC replaced cholesterol at L246 with a similar binding time was seen (Figure 5e,f). Further, L246 was found to sample two distinct conformations (Figure 5i) in the presence of 27-OHC, indicating a strong binding influence. The enhanced 27-OHC binding at N192 illustrates the C4 type interaction (Figure 5g,h with snapshots in Figure 5j). N192, an unfavourable site for cholesterol in both the CH and CH-27OHC systems is a favoured binding site for 27-OHC in the CH-27OHC system resulting in a significant conformational change. Similar analysis of the binding times and angles for other critical signalling residues in the CH-7BHC (Figure S15, C1-C4 type) and CH-27OHC (Figure S16, C1-C3 and S17, C4 type) membranes reveal that 11 out of 15 residues experienced additional oxysterol interaction (C4 type) in CH-27OHC, whereas only 1 out of 11 critical residues was influenced in CH-7BHC. The results are summarized in Table 2, where the maximum perturbation to the critical signalling residues was observed for 27-OHC due to enhanced oxysterol binding. These residues are known to be involved in signal initiation (P42, Y45, H203), chemokine engagement (N192), signal propagation (F248, W252) and microswitch activation (Y302)[28].

**Figure 5:**
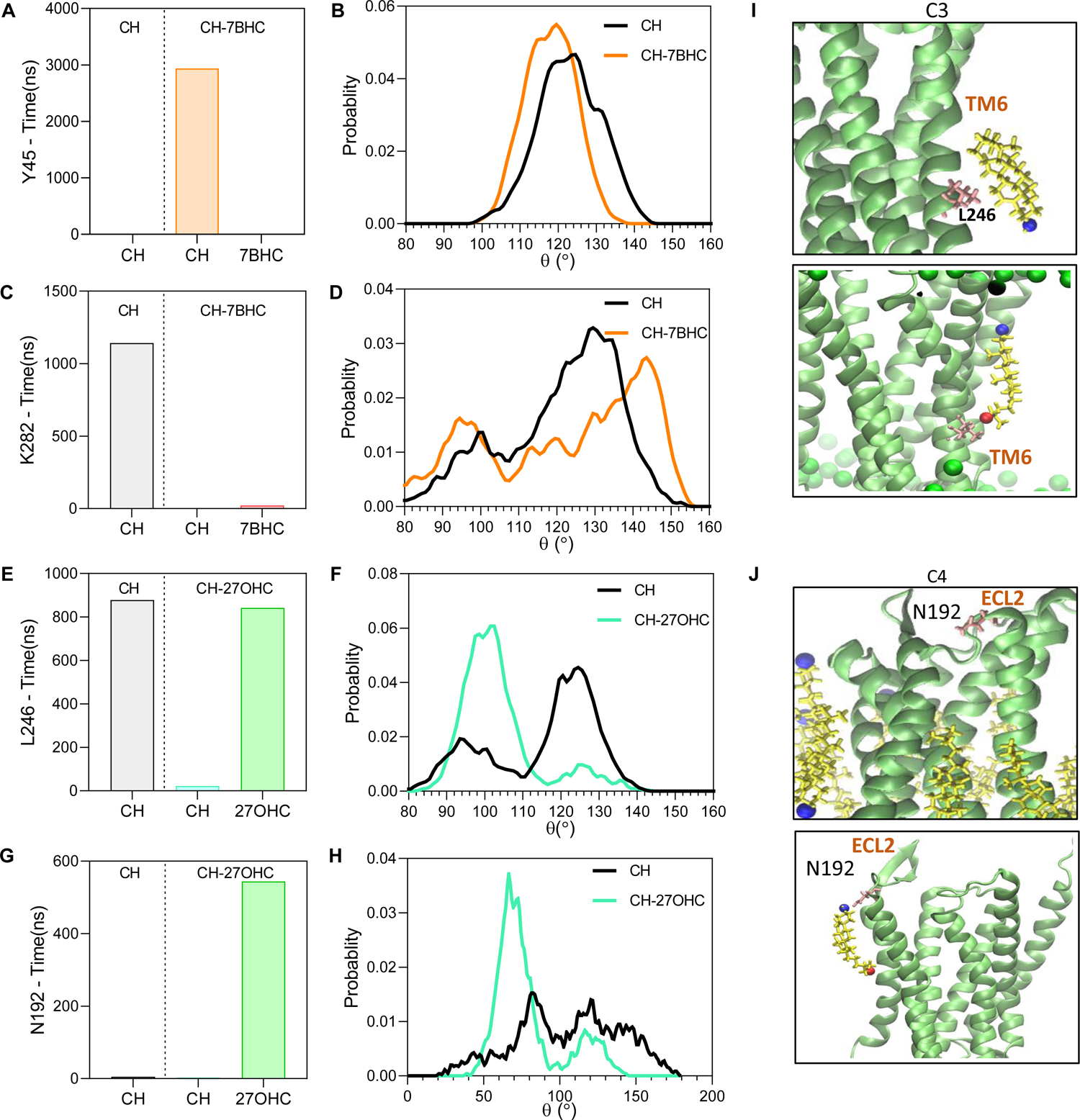
Effect of sterol interactions on critical signalling residues. An example of each of the different types of interactions (C1-C4) affecting the conformation of the critical residues, where the change in their interaction time compared to CH is shown in the left panel (a,c,e and g) and the change in θ is illustrated in the right panel (b,d,f and h). (a, b) C1-Effect of cholesterol enhancement on conformation of Y45 in CH-7BHC (c, d) C2-Effect of reduced cholesterol interaction on conformation of K282 in CH-7BHC (e, f) C3-Effect of oxysterol addition on the conformation of L246 in CH-27OHC (g, h) C4-Effect of oxysterol addition with enhanced binding on conformation of N192 in CH-27OHC. (i) Snapshots of cholesterol binding at L246 in CH (upper panel) and 27-OHC replacing it and binding to L246 in CH-27OHC (lower panel) (j) Snapshots of N192 in CH (upper panel), where none of the cholesterol molecules is binding to it, and oxysterol is shown to be interacting with it in CH-27OHC (lower panel). In snapshots, protein is shown as a cartoon in lime, P atoms of bilayer as spheres in green, sterol molecules are shown in yellow with O3 atom as a blue sphere and O27 as a red sphere.

**Table 2:**
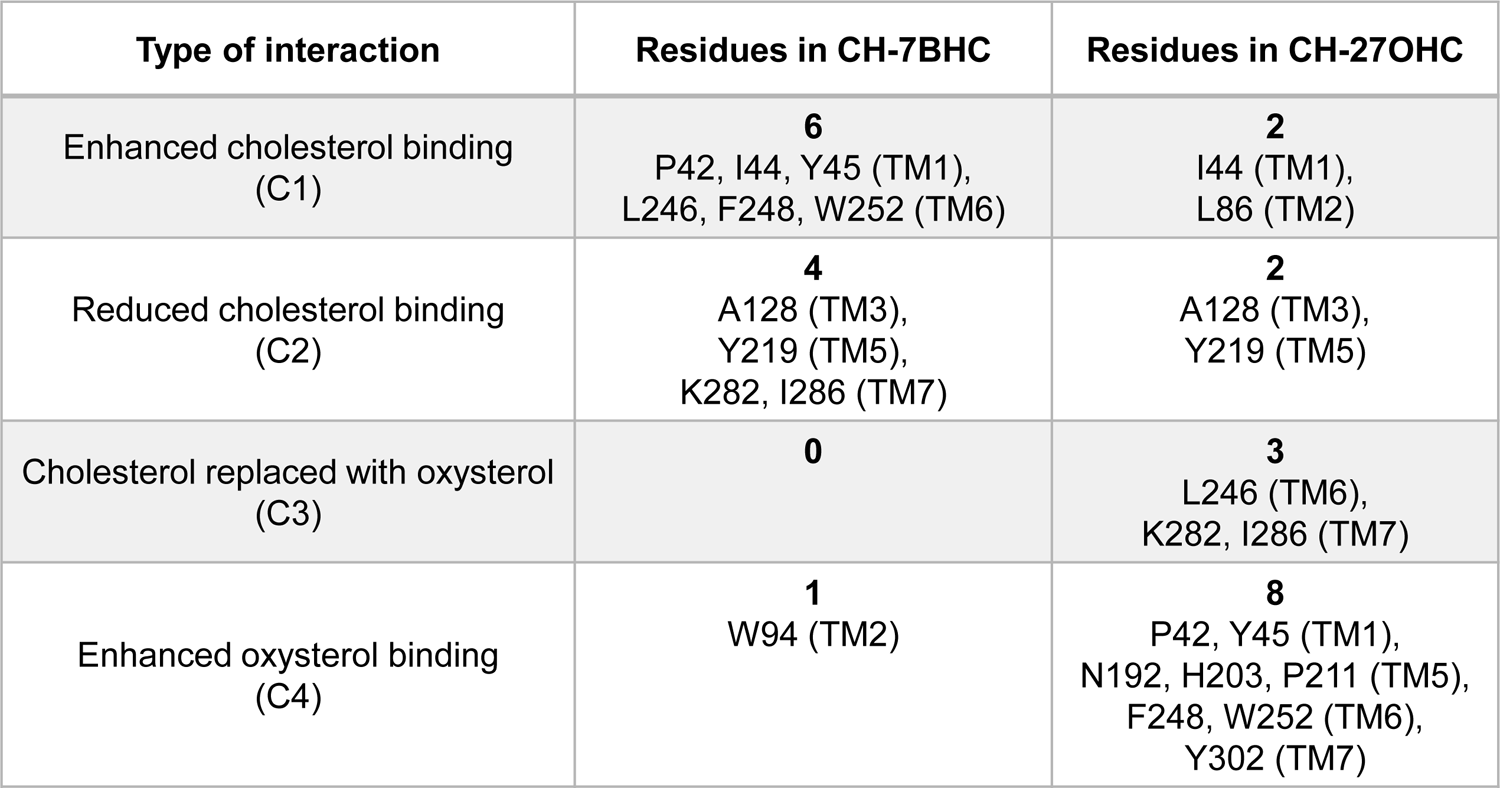
Changes in interactions of critical signalling residues in the presence of oxysterols compared with the CH-membrane. The number of affected residues is indicated in bold. See main manuscript for definition of C1-C4 type interactions. Only weak perturbations in the conformations occur at the signalling residues where C1 and C2 type interactions (Figures S15 and S16) are observed. However, for C3 and C4 type interactions where oxysterol interactions are enhanced, significant changes are observed in the conformation of the residue (Figure 5c and d main manuscript, S15 and S17). Six residues are implicated in enhanced cholesterol, C1 interactions for the CH-7BHC systems when compared with 2 residues for the CH-27OHC system. For C2 type interactions, four residues are implicated in the CH-7BHC system with 2 for CH-27OHC. The situation changes for the C3 and C4 type interactions where 3 and 8 residues are impacted in the CH-27OHC system respectively and only 1 residue is influenced in the CH-7BHC system.

These observations provided compelling evidence that the addition of 7-BHC to the plasma membrane increases or reduces cholesterol binding at the critical signalling residues in CXCR4. Similarly, the addition of 27-OHC resulted in both enhanced 27-OHC binding and cholesterol replacement, inducing significant conformational changes to the critical signalling residues. Taken together with the experimental findings, where the addition of tail oxysterols mitigates signalling to a greater extent when compared with the ring oxysterols, the loss of signalling can be attributed to the dominant effect of 27-OHC on the critical signalling residues of CXCR4.

### Sterol composition influences dimeric interface and the kink angle

Since the formation of higher-order oligomers of CXCR4 was reduced in senescent cells, we analyzed the MD trajectories to see if sterol binding at the dimeric interface was also altered. The binding time of sterols with the residues at the dimeric interface TM5[30] (Figure S18a), located in the vicinity of the extracellular surface of the receptor, revealed that binding of 27-OHC (Figure 6a) is significantly higher compared to cholesterol in the CH membrane. In sharp contrast, the binding of 7-BHC in the same region was found to be weak, with binding times below 0.5 *µs*, unlike the binding duration for 27-OHC, which was above 1.3 *µs*. This affinity for 27-OHC at the dimeric interface explains the reduced receptor clustering observed in senescent cells (Figure 3c) and cells with increased oxidized sterols (Figure S5). Further, the toggle switch, defined as the distance between Y219 and Y302, a signature for G-protein activation[31], decreased with 27-OHC (Figure S18b) to a slightly greater extent than 7-BHC (Figure 6b).

**Figure 6:**
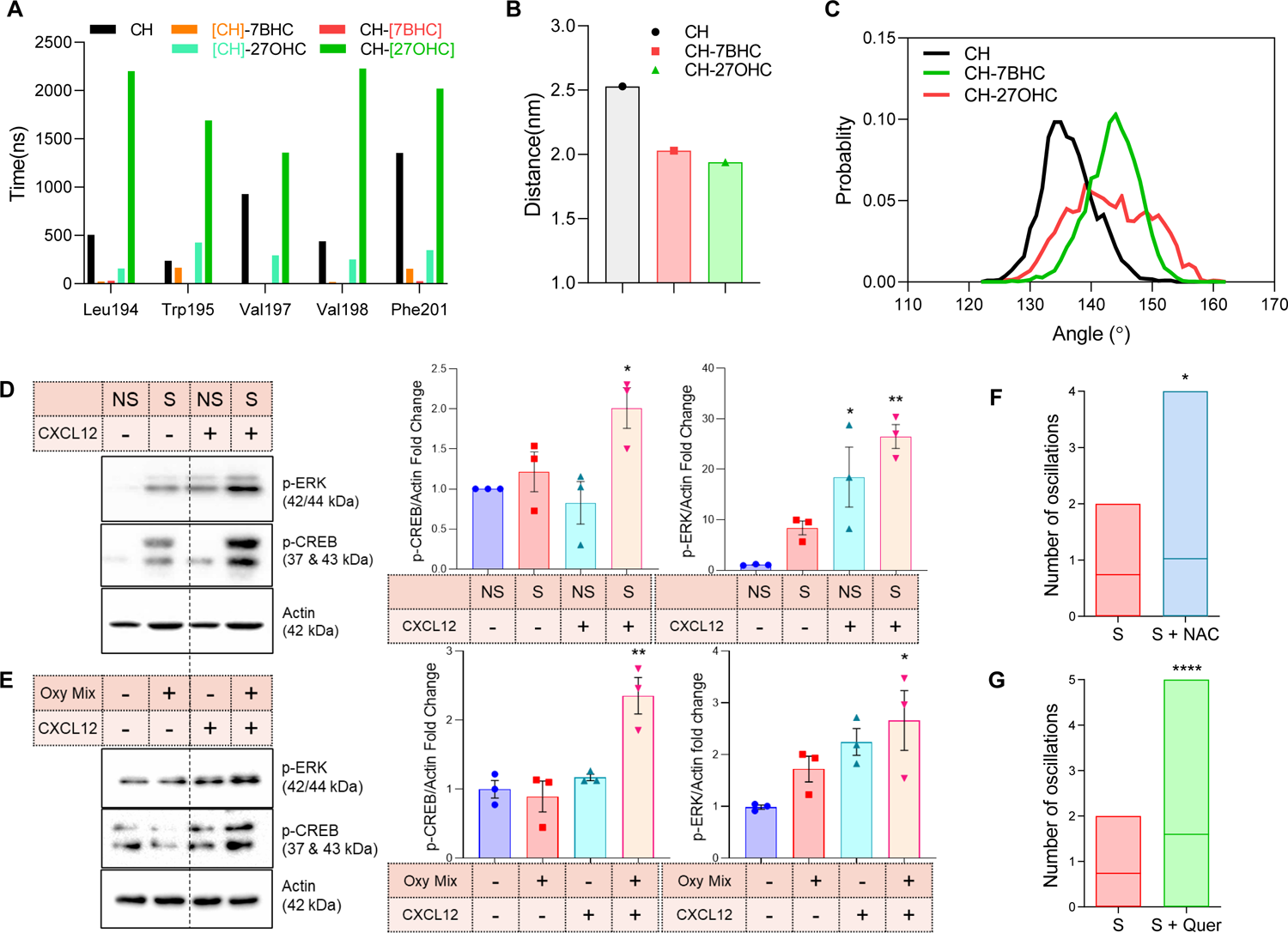
Interaction of sterols with CXCR4 alters signalling specificity. (a) Interaction time of residues present at the dimeric interface (as indicated) with various sterol molecules (b) Center-of-mass distance between the toggle switch residues, Y219 and Y302 averaged over the full trajectory (c) Kink angle distribution of TM6 calculated as the angle between Cα atoms of I245, P254 and G258 in various cholesterol-oxysterol systems. (d) Western blot analysis for p-CREB, and p-ERK in non-senescent (NS) and senescent (S) cells post stimulation (left) and its quantification (right). (e) Western blot for p-CREB and p-ERK in cells treated with oxysterol mixture post stimulation (left) and its quantification (right). Mean calcium oscillations in senescent cells treated with (f) 10 mM NAC (Movie S6) and (g) 2 μM Quercetin (Movie S7). (n=3; N > 100 for all groups).

Additionally, TM6, essential for both G-protein activation and dimerization[27, 28], possesses a characteristic kink[32], whose angle changes upon receptor activation[31]. We observed a clear shift of more than 10° in the kink angle of TM6 in CH-27OHC compared to CH (Figure 6c). Interestingly, after ligand binding, both the kink angle and toggle switch distance decrease in the active state; however, the presence of oxysterols appears to activate these signatures even in the absence of the ligand. The above signatures provide additional evidence of the interference of oxysterols in perturbing the activation of the G-protein, leading to the compromised signalling observed in senescent and oxysterol-treated cells.

These observations based on MD analysis provided reasons to investigate the impact of oxysterol on different outputs generated by coupling to different G-proteins viz. Gα_i/o_ mediated or Gα_s_ mediated-response. These changes in the output through alternation in G-protein interaction were examined by monitoring the activation status of known downstream effectors in the treated cells through western blotting. For this, we examined the activation of known CXCR4 effectors, which included CREB, a Gα_s_-cAMP-PKA dependent effector and pERK, a CXCR4 dependent MAPK transactivation effector. We recorded that CREB phosphorylation was higher in senescent cells (Figure 6d) and in oxysterol-treated non-senescent cells (Figure 6e), compared to baseline, which typically increases through Gα_s_ activation. This pointed towards an often observed but previously unexplained switching in the signalling from the Gα_i/o_ subunit, which inhibits AC and decreases cAMP levels, to Gα_s_ subunit type, which activates the cAMP-dependent PKA signalling.

Interestingly, Gα_s_ activation also affected non-canonical GPCR-MAPK signalling, and enhanced ERK phosphorylation was detected in the presence of oxysterols post-stimulation and in senescent cells where oxysterol levels were higher. These evidences demonstrated that the oxysterols accumulation in senescence causes a class switch in CXCR4 coupling from Gα_i/o_ to Gα_s_, which in turn alters the dynamics of various signalling processes during senescence. The evidence of switching comes from the suppression of calcium release, enhancement of cAMP-mediated signalling, ERK signalling and simulations for sterol occupancy on CXCR4 receptor residues. Long-term response of this switching is also evident from the absence of inflammation enhancement due to an increase in cAMP, unlike when cells are transitioning into senescence, and a reduction in cAMP is recorded, through activation of canonical Ga_i/o_ subunit. Thus, for the first time, we present the role of oxysterols in modulating GPCR signalling by regulating G-protein coupling specificity.

Based on the computational and experimental results, it was evident that the oxysterol generation in the senescent cell affects signalling through the CXCR4 receptor. The cause for oxysterol formation remains unknown; however, it is known that non-enzymatic oxidation through enhanced ROS levels in senescence could be one of the factors contributing to this process. To evaluate this aspect, we tested whether the signalling can be restored by quenching ROS using two known quenchers, N-acetyl L-Cysteine (NAC) and Quercetin. In the treated cells, there was a significant increase in the number of oscillations in cells (Figure 7f, g), and partial restoration of signalling was recorded. Overall, our studies demonstrate a crucial missing link between dysregulated GPCR signalling in aging and show how elevated oxidative stress in the aged can alter the signalling through various membrane-anchored proteins through modulation of the chemical composition of membrane-cholesterol.

**Figure 7:**
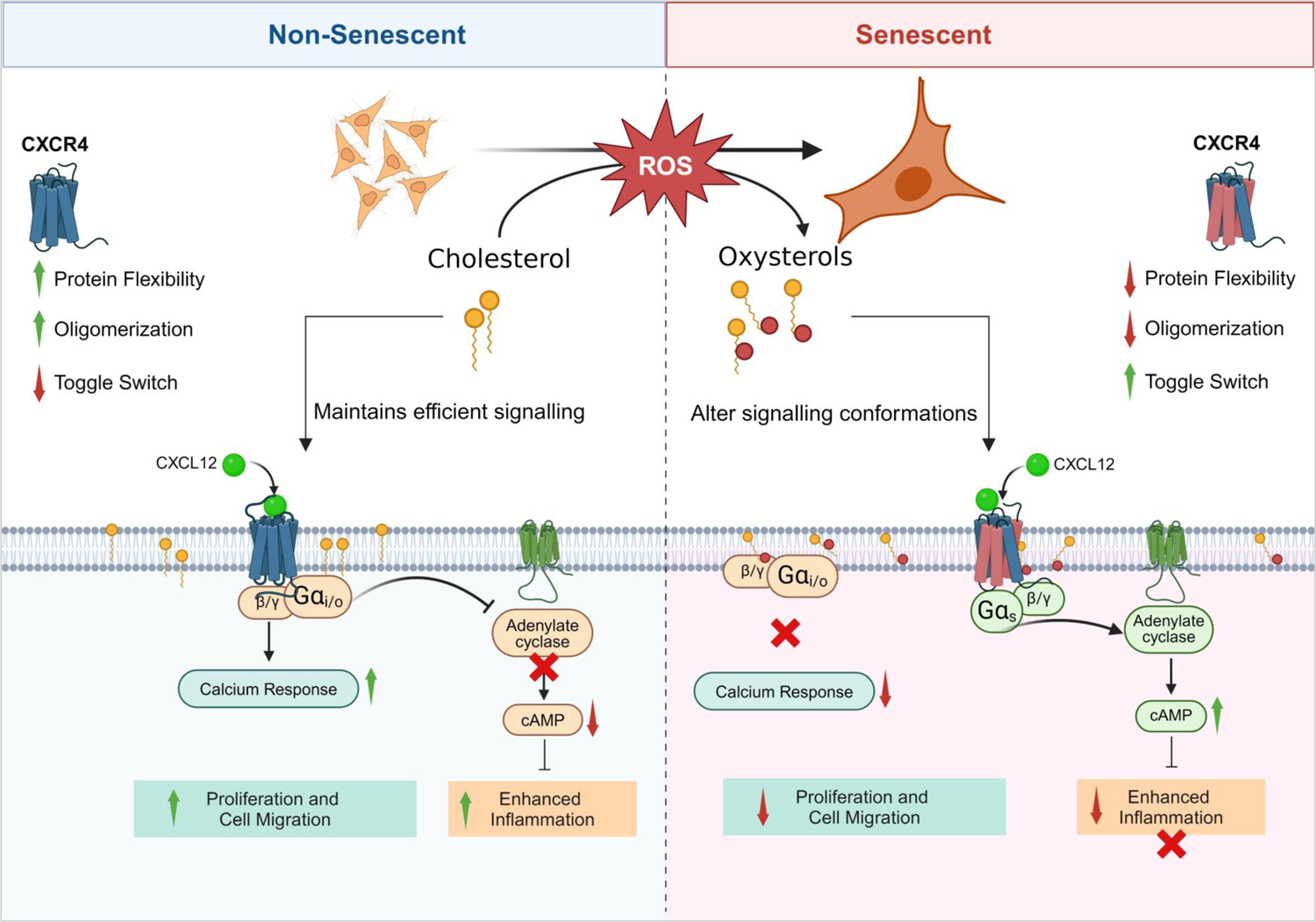
Model of findings from computational and experimental studies for altered CXCL12-CXCR4 signalling in senescent cells. This study identified oxysterols as modulators of GPCR signalling during senescence for the first time. The chemokine receptor CXCR4 is upregulated in senescent cells, however, the CXCL12-mediated calcium response is impaired, and the downstream signalling is altered. Cholesterol upon oxidation generates oxysterols which can disrupt membrane properties. Our experimental evidence demonstrate that senescent cells behave like a cholesterol-depleted or oxysterol-enriched environment. The effect of this alteration in cholesterol-oxysterol ratio was captured on CXCR4 receptor organization by MD analysis and validated by experimental approaches. Our study reveals that the CXCR4 receptor signals through the Gαi/o G-protein; however, the presence of oxysterols switches the affinity towards Gαs. This switching converts inflammatory signalling from a pro-inflammatory pathway to an anti-inflammatory pathway. The CXCR4 signalling was partially restored in the presence of ROS quenchers, which can potentially prevent cholesterol to oxysterol conversion. Overall, we reveal an unanticipated role for oxysterols in aging and explain refractory behaviour of receptors in aged cells.

## Discussion

Our study explains how a GPCR, CXCR4 receptor interacts with cholesterol and oxysterols and how these interactions affect signalling outcomes during cellular aging (Figure 7). While the role of cholesterol modulating GPCR activity is well established, this is the first report in which the physiological oxidation product of cholesterol, called oxysterol, modulates receptor signalling outcomes in a physiological condition, aging. Previously, the roles of CXCR4 in aging, inflammation and cancer metastasis have been reported. However, many instances are reported in literature where CXCR4 couples to Gα subunits other than its canonical Gα_i/o_ subunits[33, 34] or lead to differential signalling outputs[35, 36], and the exact mechanism or conditions under which this occurs remained unexplored.

Our study provides the first direct evidence of oxysterol accumulation in physiologically relevant senescent conditions and how they influence signalling outcomes by modulating G protein class switching. We show that senescence is associated with high reactive oxygen species and upregulation of cholesterol biosynthetic pathways. These independent observations led us to investigate the oxidation of cholesterol in senescent cells. We report that oxysterol accumulation impairs CXCR4 receptor calcium signalling, a ubiquitously expressed GPCR essential for various cellular processes, including cell division, immune cell migration and inflammation. The impaired G-protein signalling in senescent cells could have several negative consequences, including reduced immune cell function and migration with age. In neutrophils, the CXCR4 receptor is responsible for homing senescent neutrophils to the bone marrow through a chemotactic response to the stromal gradient of SDF1α[37]. Incidentally, it has been seen that aged mice have an abundance of senescent neutrophils in circulation[6], which could now be attributed to oxysterol accumulation, which leads to loss of CXCR4 receptor-dependent chemotactic function. It could also lead to impaired cell proliferation and differentiation, and it is well known that senescent cells are refractory to mitogenic stimuli[38]. The presence of oxysterols in senescent cells thus can explain the loss of response in these cells, which could contribute to tissue aging and dysfunction.

Our findings emphasize the central and modulatory role played by membrane sterol constituents on CXCR4 signalling. The presence of tail oxysterols showed impaired calcium response when compared with the ring oxysterols. In the case of 22R-OHC, calcium signalling was completely abolished, followed by 27-OHC and 25-OHC. All-atom MD simulations with cholesterol and cholesterol mixtures with 7-BHC and 27-OHC provide several molecular insights to decipher the modulated signalling outcomes observed in the experiments. Dramatic reduction in flexibility of CXCR4 was revealed in the RMSFs in the presence of membrane oxysterols suggesting that flexibility in the native state could potentially compromise conformational changes required for signal transduction as seen in other membrane-protein systems[39, 40]. The most striking effects were observed with 27-OHC, where a near-complete replacement of cholesterol was observed at several residues implicated in signal propagation and initiation.

The displacement of cholesterol by 27-OHC was the greatest in the transmembrane motifs TM5, TM6 and TM7 implicated in signal activation and propagation[27] as well as at residue Y302 implicated in microswitch activation. These results illustrate the manner in which critical signalling residues are influenced by the presence of oxysterols with a clear distinction observed between the ring and tail oxysterols. This difference in oxysterol binding provides a direct connection with the greater reduction in calcium oscillations observed in the experiments with the tail oxysterols when compared with the ring oxysterols. Clearly, the enhanced binding of 27-OHC with the critical signalling residues coupled with their reorientation, compromises signal transduction and activation of CXCR4 as observed in the mitigated signalling in the presence of membrane oxysterols.

Our findings are also consistent with the existing knowledge that oxysterols play a role in the development of age-related diseases[41]. The role of 7-KC has been most widely studied and implicated in age-related pathological conditions[42]. Several studies have reported the involvement of oxysterols in pathological conditions like neurodegenerative diseases, cancers and metabolic disorders[43–45]. For example, oxysterols have been shown to accumulate in the brains of patients with Alzheimer’s disease[46] and in the atherosclerotic plaques of patients with cardiovascular disease[47]. Oxysterols have also been shown to promote tumor growth and progression[48].

Given the effect of oxysterols on GPCR signalling, a novel mechanism of dysregulation of signalling in pathological conditions can be explained by our combined experimental and molecular dynamics study. G-protein class cross-coupling has been demonstrated for several GPCRs[49], affecting processes like cell-fate decision, survival, migration and proliferation. We show that oxysterols cause a class switch in CXCR4 coupling from Gα_i/o_ to Gα_s_ and based on this, a detailed analysis of G-protein coupling with CXCR4 and other GPCRs in the presence of oxysterols needs to be investigated to understand the molecular mechanisms involved in this switch. Furthermore, macroscopic kinetic models have been used to study calcium oscillations[50] and similar models will be needed to understand the loss of signalling and class switching in the presence of membrane oxysterols. Overall, we present an alternate paradigm in GPCR signalling, wherein GPCR-G-protein coupling specificity is altered by the levels of oxysterols in cells. Given that oxysterol levels are intricately associated with the levels of reactive oxygen species in a cell, which in turn is linked to metabolic processes, our study paves the way for metabolic remodeling of signalling for GPCRs, an area which needs detailed investigations and may help us address the changes in the efficacy of drugs used to treat aging populations.

## Methods

### Cell culture and treatments

HeLa, Beas2b, HaCat and LX2 cell lines were obtained from ATCC and cultured in DMEM (Sigma) containing sodium bicarbonate 3.7 g/L, sodium pyruvate 0.11 g/L, penicillin and streptomycin 100U/ml and 10% FBS. Cells were cultured at 37°C in the presence of 5% CO_2_ in a humidified incubator. Senescence was induced by treating HeLa cells with 200μM 5-Bromo-2’-deoxyuridine (BrdU, Sigma) freshly prepared in DMSO or exposing the cells to ionizing radiation (8 Gy) and incubation for 4 days. Primary cells were exposed to ionizing radiation (8 Gy) for inducing senescence. Beas2b and HaCat were incubated for 4 days and LX2 for 6 days to induce senescence. For cholesterol depletion, 1×10^5^ cells were seeded in a 35 mm glass-bottom dish and allowed to adhere overnight. Cells were treated with 2.5, 5 or 10 mM methyl-β-cyclodextrin (mβCD) (Sigma-Aldrich) in DMEM for 1 hour at 37°C. For oxysterol treatments, cells were treated with 10 µg/mL of each oxysterol (27-OHC, 25-OHC, 22(R)-OHC, 24(S)-OHC, 7-KC or 7-BHC) (Cayman Chemicals) or 10 µL ethanol (vehicle) in DMEM for 1 hour at 37°C. Oxysterol mixture treatment was done by adding each oxysterol to a total final concentration of 10 µg/mL in DMEM for 1 hour at 37°C. For quenching ROS, cells were treated with 10mM of NAC (Sigma) or 2µM Quercetin (Sigma) for 24 hours in complete medium.

### SA-β-Galactosidase Staining

The protocol described in Dimri *et al*.[51], was followed. Briefly, cells were washed with 1x phosphate buffer saline (PBS) twice and fixed with 0.2% glutaraldehyde for 15 minutes at room temperature (RT). Cells were washed thrice with 1x PBS followed by the addition of staining solution (40mM citric acid/sodium phosphate buffer, 5mM potassium ferrocyanide, 5mM potassium ferricyanide, 150mM sodium chloride, 2mM magnesium chloride, X-gal 1mg/ml). Cells were incubated overnight at 37°C, followed by removal of the staining solution, washing thrice with 1x PBS, and images were acquired using an inverted epifluorescence microscope (Olympus IX81, Japan) with a 20X objective.

### Surface staining

Cells were seeded in 6-well plates and allowed to adhere overnight, followed by treatments. Cells were washed 3 times with 1x PBS, fixed in 4% paraformaldehyde for 10 mins at RT. Cells were washed, followed by blocking using 10% FBS in PBS for 30 mins. The primary antibody was diluted in antibody dilution buffer and incubated for 30 mins at RT, followed by 1x PBS wash. A secondary antibody conjugated with Alexa-488 (Thermofisher) was incubated for 30 mins at RT in the dark, followed by 1x PBS wash. Surface expression analysis was done by imaging using an inverted epifluorescence microscope (Olympus IX83, Japan) with a 60X objective.

### ELISA

Cells were seeded in 24-well plates and allowed to adhere overnight, followed by addition of 100 ng/mL CXCL12 and the supernatant was collected 48 hours post-treatment. Per the manufacturer’s instructions, the levels of secreted cytokines were quantified using the BD OptEIA™ Human IL-6 ELISA kit (BD Biosciences). Cells were trypsinized and counted for normalization of cytokine levels per cell.

### Gene Expression Profiling

#### RNA Isolation

To isolate total RNA, 2×10^5^ cells were seeded in 6-well plates followed by treatments. For cell lysis, 1 ml of RNA Iso Plus (Takara Bio) was added to cells and incubated for 5 minutes at RT. The cell lysate was collected, and RNA extraction was performed using the chloroform-isopropanol extraction method. 200mL chloroform was added, followed by vigorous shaking and incubation for 10 mins at RT. Samples were centrifuged at 12000g for 15 mins at 4°C, and aqueous phase supernatant was collected. 500 mL isopropanol was added to the supernatant and centrifuged at 12000g for 10 mins at 4°C to obtain RNA pellet. The pellet was washed with 75% ethanol and centrifuged at 7500g for 5 mins at 4°C. The RNA pellet was resuspended in nuclease-free water, quantified using ND-1000 UV-Vis Spectrophotometer (NanoDrop Technologies), and stored at −80°C.

#### cDNA synthesis

cDNA was synthesized using the Bio-Rad iScript^TM^ cDNA synthesis kit as per the manufacturer’s instructions. Briefly, 1mg of RNA was used for the reaction described below, and the cDNA was used for qPCR analysis.

| Component | Volume per reaction |
| --- | --- |
| 5x iScript Reaction Mix | 4 mL |
| iScript Reverse Transcriptase | 1 mL |
| Nuclease Free water | Variable |
| RNA (1 mg) | Variable |
| **Total** | **20 mL** |

#### qRT-PCR

Real-Time PCR reaction was set up using 25ng of cDNA and DyNamo Flash SYBR Green qPCR kit (ThermoFischer Scientific) according to the manufacturer’s protocol. Acquisition and data analysis were done using Rotor-Gene Q (Qiagen) qPCR machine and accompanying software. Primer sequences used for gene expression profiling are:

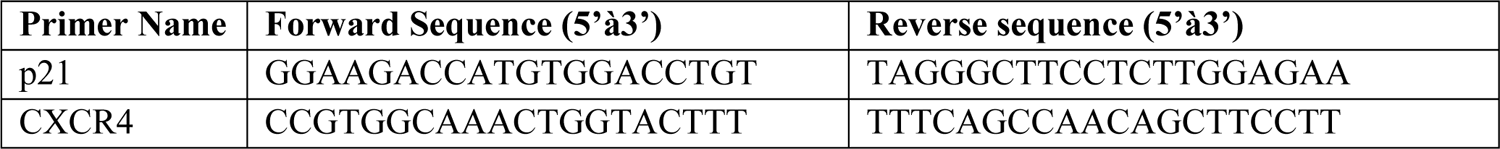

### Microarray analysis

For microarray analysis, Agilent Custom Human Gene expression 8X60K microarrays were used. Spot intensity data was deposited in Gene Expression Omnibus (GEO) with accession ID: GSE254769. Gene expression data contained information of four groups: non-senescent, senescent, non-senescent with CXCL12 stimulated and senescent CXCL12 stimulated samples of HeLa cell line. Each group consisted of two replicates. The response index (RI) was calculated for individual genes as follows:

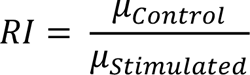

where μ represented the average of the two replicates’ spot intensity measurements. After calculating the ratio between the log2-transformed response index of senescent (RIS) and response index of non-senescent (RINS), transcripts with RINS and the ratio value threshold of ±0.05 were taken into consideration for further analysis. Each group’s average spot intensity value was divided by non-senescent and log2-transformed values were considered for analysis. Heatmap was created using Pheatmap package[52] in R 4.3.2. For RIS and RINS heatmap, z-score transformed values were taken. For Gene Set Enrichment Analysis (GSEA), BioMart on Ensembl changed transcript symbols to Ensembl IDs[53, 54], and the clusterProfiler package[55] was used to perform GSEA on the available Ensembl IDs. GSEA was performed for each of the three aspects (biological process, cellular component, and molecular function) and the cellular component using the Gene Ontology (GO) gene set and top 20 most enriched and significant pathways were plotted. The GO chord plot was made using SRplot web browser for selected GO terms of RIS and RINS ratio[56].

### DCFDA for ROS measurement

Cells were seeded in a 24-well plate and incubated overnight for adhesion to detect intracellular ROS. Cells were washed with 1x PBS and incubated with 10μM of 2’, 7’ – dichlorofluorescein (DCFDA) (Sigma Aldrich, USA) in DMEM for 30 minutes at 37°C in the dark. Cells were washed thrice with 1x PBS, and fluorescence intensity was recorded using the Infinite M1000 plate reader (Tecan) at 492/525 nm excitation/emission. Cells were trypsinised and counted for normalisation of fluorescence intensities to cell number and the ROS levels per 1000 cells were plotted.

### Fluorescence recovery after photobleaching (FRAP)

Cells stably expressing CXCR4-GFP were seeded in a glass bottom dish, followed by various treatments. Live cell imaging was done using an Olympus IX83 epifluorescence microscope with a stage-top Uno CO_2_ incubation system (OKOLab) at 37°C. The selected region of interest on the cell membrane was photobleached at the 5^th^ time point using a 488 nm laser for 300ms at 100% power using the 3i vector photomanipulation system (3i, USA), controlled by Slidebook 6.0 Software. Fluorescence recovery was measured using time-lapse imaging in the GFP channel for 15 minutes at a 15-second interval between two successive frames. For calculating fluorescence intensity at each time point, the intensity of the bleached regions was normalized to maximum pre-bleach intensity. The relative intensity was then plotted as (I_t_ - I_o_), where I_t_ refers to the normalized intensity at the time point post-bleaching under analysis, and I_o_ is the normalized intensity pre-bleaching, as a function of time. Non-linear curve fitting of the resultant plot was used to calculate the t_1/2,_ which indicated the recovery time.

### FLIM

HeLa cells stably expressing CXCR4-GFP were seeded at a density of 1×10^5^ cells per dish, followed by senescence induction or oxysterol treatment. The cells were washed twice with 1x PBS and imaged in DMEM without phenol red. A Leica TCS SP8 X microscope with 63x oil immersion objective was used for FLIM. The sample was excited at a wavelength of 488 nm with a laser intensity set at 30%. Emitted light was collected using the green channel at 580 nm, extending up to approximately 12ns or until 100 photons were acquired. Analysis of the acquired data was done using Leica LAS-X software. Five regions of interest were randomly selected on each cell membrane within the frame to determine their fluorescence lifetime values. The decay curve was fitted according to an exponential model, with the selection of the best-fit curve based on the minimal residues. For optimal fit, the lifetime values with a χ^2^ between 0.99 and 1.10 were taken for statistical analysis. The mean lifetime was obtained from the range of lifetimes obtained across each treatment and plotted.

### Receptor Internalization

HeLa cells stably expressing CXCR4-GFP were seeded in 35mm glass bottom dishes followed by oxysterol treatments, or BrdU treated senescent cells expressing CXCR4-GFP were seeded. Cells were washed with 1x PBS, and an imaging medium was added. Imaging was done using Olympus FV10i confocal laser scanning microscope. CXCL12 was added and imaged for 30 mins to record receptor internalization. Percentage internalization was plotted using a ratio of the cytoplasmic by whole-cell intensity.

### Oxysterol Estimation

#### Lipid Isolation

1×10^6^ cells (HeLa, Beas2b, HaCat and LX2) were seeded in 100mm cell culture dishes followed by treatments. Cells were washed with 1x PBS and centrifuged to collect the cell pellet. Total lipid extraction was done by adding 200 mL of chloroform-methanol (v/v 2:1) and centrifugation at 13000rpm for 10 mins at RT. The supernatant containing lipids was collected into a fresh LC-MS glass vial (Agilent Technologies), and the organic solvent was vacuum dried. The lipids were dissolved in ethanol, and the sample was directly processed for LC-MS.

#### Oxysterol Derivatization

The oxysterol standards and samples were derivatized using the Oxysterol Derivatization MaxSpec Kit® (Cayman Chemicals) according to the manufacturer’s instructions. For cholesterol, 6.25µg/mL to 100 µg/mL concentration points of standard curve whereas for its derivatives 7-BHC, 22(R)-OHC, 24(S)-OHC, 25-OHC and 27-OHC 0.62ng/mL to 20 ng/mL concentration points were made. Biological samples were acquired twice (undiluted and 10 times dilution). The standard for 7-KC was not detected in our protocol, hence its quantification was not performed.

#### LC-ESI-MS/MS

Samples were acquired on Exion AD UHPLC chromatography system coupled with ZenoTOF 7600 system (SCIEX) and were ionized using electrospray ionization (ESI) source. Chromatography separation was achieved by utilizing F5 column (100 x 3mm, 2.6µm) with a column oven of 40°C with injection volume 30 µL. Gradient elution was used with a flow rate of 300 µL/min. Mobile phase A and B used for separation consisted of water with 0.1% formic acid and acetonitrile with 0.1% formic acid, respectively. Chromatographic gradient used for separation is as follows: the initial mobile phase was started at 0% B followed by 15% B from 0–1 min, 80% B from 1–10 min, 95% B from 10 –10.2 min; kept at 95% B from 10.2-13 min, 2% B from 13–13.1 followed by equilibration at 2% B from 13.1–16 min with a flow rate of 300 µL/min.

LC-MS acquisition was performed in MRMHR mode by utilizing electron-activated dissociation (EAD) for fragmentation in positive ionization. The parameters used for mass spectrometry are as follows: nebulizer gas (GS1) = 35 psi; desolvation gas (GS2) 40 psi; curtain gas flow = 30 psi; capillary voltages (ISVF) = 5500 V; source temperature = 400°C.

For TOF MS, mass range was set to 350 to 650 Da with an accumulation time of 0.25 seconds, declustering potential (DP) of 80 V and collision energy of 10 V. Whereas for MRM-HR, m/z 539.44 was selected for EAD fragmentation with electron kinetic energy of 23, accumulation time 0.2 seconds and a mass range of 50-600 Da.

For quantitation of cholesterol and oxysterols, unique fragments were selected for 27-OHC, 25-OHC, 24(S)-OHC, 22(R)-OHC whereas for 7-BHC and cholesterol parent mass was selected for quantitation (Figure S4a). The table below shows corresponding retention time and selected fragment used for quantitation for cholesterol and oxysterols.

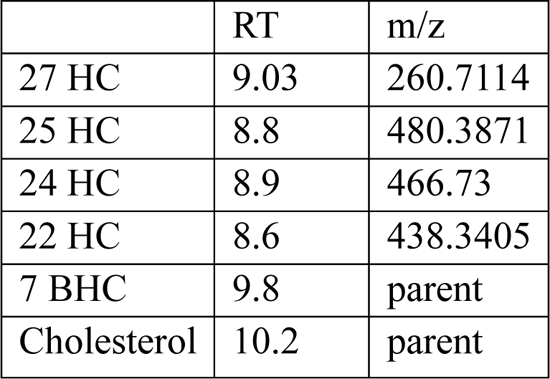

|  | RT | m/z |
| --- | --- | --- |
| 27 HC | 9.03 | 260.7114 |
| 25 HC | 8.8 | 480.3871 |
| 24 HC | 8.9 | 466.73 |
| 22 HC | 8.6 | 438.3405 |
| 7 BHC | 9.8 | parent |
| Cholesterol | 10.2 | parent |

***Sample processing*** was performed using Analytics in SCIEX OS software.

### Resazurin Assay

Cells were seeded in a 24-well plate, followed by treatments. Cells were washed with 1x PBS, and Resazurin dye was added at a concentration of 10 μg/mL in DMEM. Cells were incubated for 3-5 hours at 37°C in the CO_2_ incubator. After incubation, fluorescence was recorded using the Infinite Pro M1000 plate reader (Tecan) at 560/590 nm excitation and emission wavelengths.

### Filipin Staining

The staining was performed using the cholesterol estimation kit (Cayman Chemicals) per the manufacturer’s protocol. In a 6-well plate, 2×10^5^ cells were seeded overnight, followed by washing with 1x PBS and treatments. Cells were washed with 1x PBS, trypsinized, and fixed with 3% paraformaldehyde for 1 hour at RT. Cells were washed with 1x PBS and incubated with 1.5 mg/ml glycine in PBS for 10 minutes at RT to quench paraformaldehyde. 0.05 mg/ml Filipin staining solution in PBS was added to cells and incubated for 2 hours at RT in the dark. Cells were washed with 1x PBS, followed by flow cytometry using BD Influx^TM^ Cell Sorter by UV excitation at 355 nm and recording emission at 480 nm.

### Calcium release assay

1×10^5^ cells were seeded in a glass-bottom dish and treated with various agents as indicated, followed by washing with HBSS (without calcium and magnesium). The calcium-binding dye Fluo-4AM (F14201, ThermoFisher Scientific, USA) was added at a concentration of 1M in HBSS and incubated for 30 minutes at 37°C. The cells were washed with HBSS and imaged in HBSS containing 10 mM HEPES buffer (pH 7.0). Live cell imaging was done using an Olympus IX83 epifluorescence microscope with a stage top Uno CO_2_ incubation system (OKOLab) at 37°C. 20X objective was used for all calcium release assays. Cells were imaged for 2 minutes with images recorded every 1 second. 10 ng/ml CXCL12 was added at the 10th frame. Image analysis was done by measuring the intensity of the region of interest marked as whole cells in a single frame of multiple experiments and calculating relative intensity (F*_t_*/F_0_) where F_0_ is the intensity at the first frame and F*_t_* is the frame intensity at each time point before and after stimulation. Calcium peaks, duration and amplitude of the response were analyzed using MATLAB. Response time is calculated by the appearance of the first peak after stimulation. For peak duration, the full width at half maxima was calculated and per cell average peak width was plotted. The amplitude of response per cell was plotted for the first peak normalized to baseline. Movies of various experiments done for calcium release can be accessed in supplementary data files.

### Western blotting analysis

2×10^5^ cells were seeded in 60mm dishes and were starved for 24 hours before treatment with vehicle or oxysterols. Treatment was done for 1 hour followed by 30 mins of stimulation with CXCL12 (10ng/ml). Cells were lysed using RIPA buffer containing protease and phosphatase inhibitor cocktail (Cayman Chemicals). After protein quantification through BCA, 50-100mg of protein was loaded and resolved by SDS PAGE and transferred onto PVDF membrane (BioRad). The membrane was blocked using 5% BSA in TBST for 1 hour at room temperature and incubated with primary antibody overnight at 4°C. The membrane was washed 3 times with TBST followed by secondary antibody incubation for 1 hour at room temperature. The membrane was washed and developed using ECL kit (BioRad). Image analysis and quantification were performed using ImageLab software (BioRad).

### CXCR4 structure

MD simulations were carried out using the structure of the monomer CXCR4 in a membrane, using the PDB entry-3ODU. The TM prediction analysis was done (Figure S20) and the missing 51 residues were modeled using I-TASSER, and the normalized Z-score[57, 58] of 5.23 with PDB entry-3ODU suggests that the model is close to the original crystal structure. Additionally, the B-factor is less than 0 for most of the structure, indicating that the model is stable (Figure S21b).

### Membrane simulations

In addition to cholesterol (chol), we used the oxysterols, 7-*β*-hydroxycholesterol (7-BHC) and 27-hydroxy-cholesterol (27-OHC). The atomic structure of all three sterols is shown in Figure S2 of SI. The CHARMM36 force field for both the oxysterol molecules was modified with charges adjusted using the procedure by Vanommeslaeghe et al.[59], and missing dihedrals and angles were included as described in the SI. All-atom membrane MD simulations with sterols and 1-palmitoyl-2-oleoyl-sn-glycerol-3-phosphocholine (POPC) were carried out to check the validity of the force fields (Figure S22).

### CXCR4 in the membrane simulations

All-atom MD simulations were performed with CXCR4 in a membrane of POPC with 20% sterol concentration (Table 1) using the CHARMM36 force field[60, 61] and the modified TIP3P water model. Na^+^ and Cl^−^ counterions were added to maintain a concentration of 150 mM. The initial structure consisting of CXCR4 in the membrane made up of 160 POPC and 40 (20 %) cholesterol molecules, was prepared using CHARMM-GUI[62] with a prescribed orientation of CXCR4[27]. Membranes with oxysterols were prepared by replacing half the cholesterol molecules in each leaflet with either 7-BHC or 27-OHC (Table 1). The initial structures were subjected to energy minimization using the steepest-descent method with a maximum force of 1000 kJ mol^−1^ nm^−1^. The energy minimized structure was subjected to NVT (canonical ensemble) equilibration for 1 ns, followed by NPT equilibration for 80 ns. The final equilibrated structure was used as the starting structure for production runs of 3 µs each. All the simulations were performed in the NPT ensemble with GROMACS version 5.1.4[63]. Periodic boundary conditions were applied in all three directions, and the temperature was maintained at 303.15 K using a Nosé–Hoover thermostat[64] with protein, membrane, and solution coupled separately with a coupling constant of 1.0 ps. Semi-isotropic pressure control was achieved using Parinello-Rahman barostat[65] with a time constant of 5.0 ps. Long-range electrostatic interactions were calculated using the particle-mesh Ewald (PME) method[66], and hydrogen bonds were constrained using the linear constraint solver (LINCS) algorithm[67]. The pressure was kept constant at 1 bar using the isothermal compressibilities of *K_xy_* = *K_z_*= 4.5 × 10^−5^ bar^−1^.

### Binding Calculations

To evaluate the binding of sterols with CXCR4, the coordination number of a given sterol atom (i) was calculated with each protein atom (j) at time frames separated by 100 ps using the following coordination function[68]

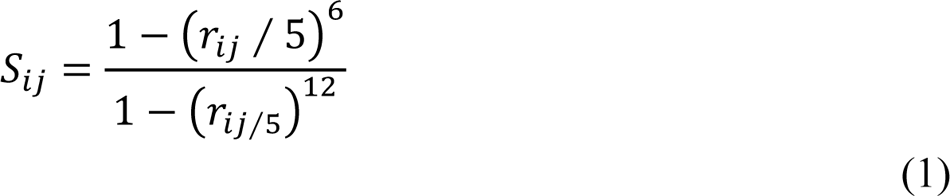

where *r_ij_* is the distance in Angstroms between the given sterol atom (*i*) and the protein atom (*j*). The value of the function is close to 1 when *r_ij_* is less than 5 A^°^ and sharply goes to zero if *r_ij_* is greater than 5 A^°^. Using the function *S_ij_*, the coordination number for each protein atom *j*, is calculated using,

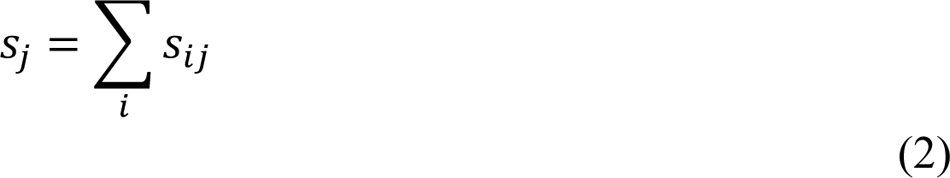

Since binding was evaluated residue-wise, we calculated the mean-coordination number of a residue *R* using,

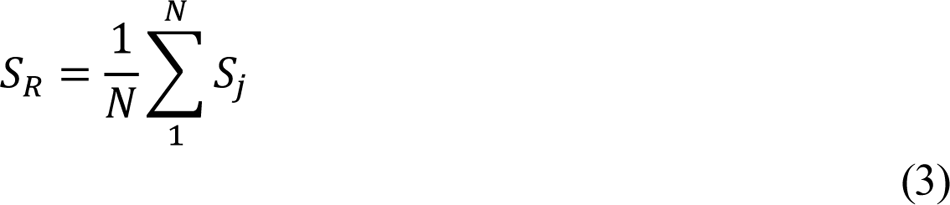

where *N* is the number of atoms in a given residue, *R*. If the *S_R_* ≥ 2 in a time frame, we assume that the sterol molecule is bound to the residue *R*. The same coordination function and criteria were also used to examine the interactions between the sterol molecules in all three membrane environments studied.

### Statistical analysis

Biological triplicates or more were used for all experiments, and the results are represented as mean ± s.e.m. The number of biological replicates (n) and the number of cells (N) analyzed are mentioned in the figure legends. Outlier removal (10% stringency) was done for all single-cell analysis data before plotting. For statistical analysis, One-way ANOVA (multiple comparisons with Untreated or Vehicle) or the Mann-Whitney test was used. Significance (p-value) is represented as *, where * ≤ 0.05, ** ≤ 0.01, *** ≤ 0.001 and **** ≤ and ns, where *>* 0.05 for ‘not significant’.

## Acknowledgments & Funding Sources

We thank the Sciex team at Bangalore for helping us develop the oxysterol estimation pipeline. We acknowledge the Thematic Unit of Excellence on Computational Materials Science (TUE-CMS) supported by the Department of Science and Technology (DST) for computational facilities. We would like to thank the Supercomputer Education and Research Center (SERC) for availing computational facility at the Indian Institute of Science, Bangalore. The funding for this work has been provided by grants from the Science and Engineering Research Board, DST, and Prashanth Prakash Family Foundation.

## Supplemental Figures

**Figure S1:**
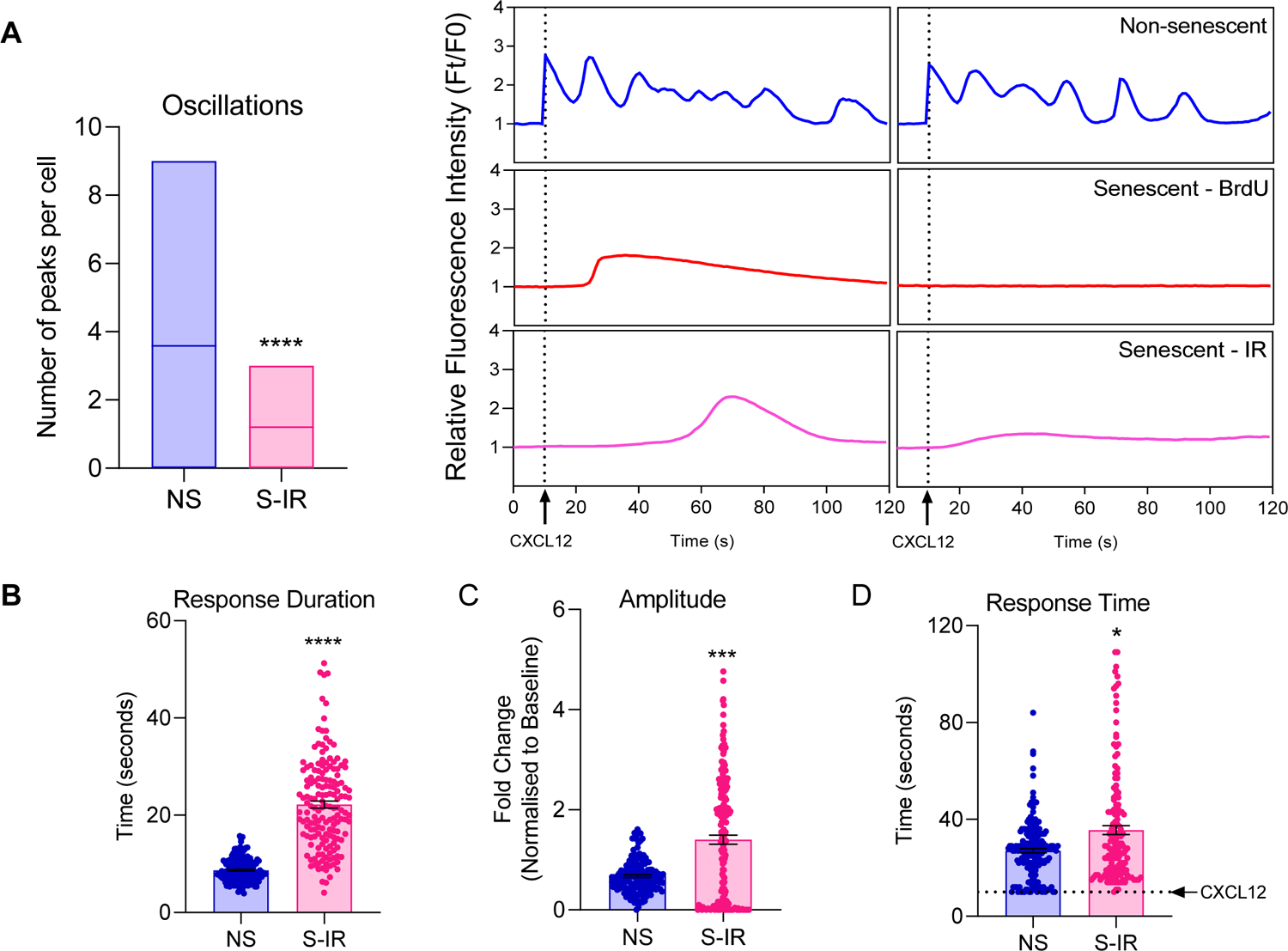
Calcium response in ionizing radiation model of senescence. (a) Mean number of calcium oscillations (left panel) and two single cell representatives (left panel). Single cell analysis of (b) response duration, (c) amplitude and (d) response time in responding cells (n=3; N>100 for all groups).

**Figure S2:**
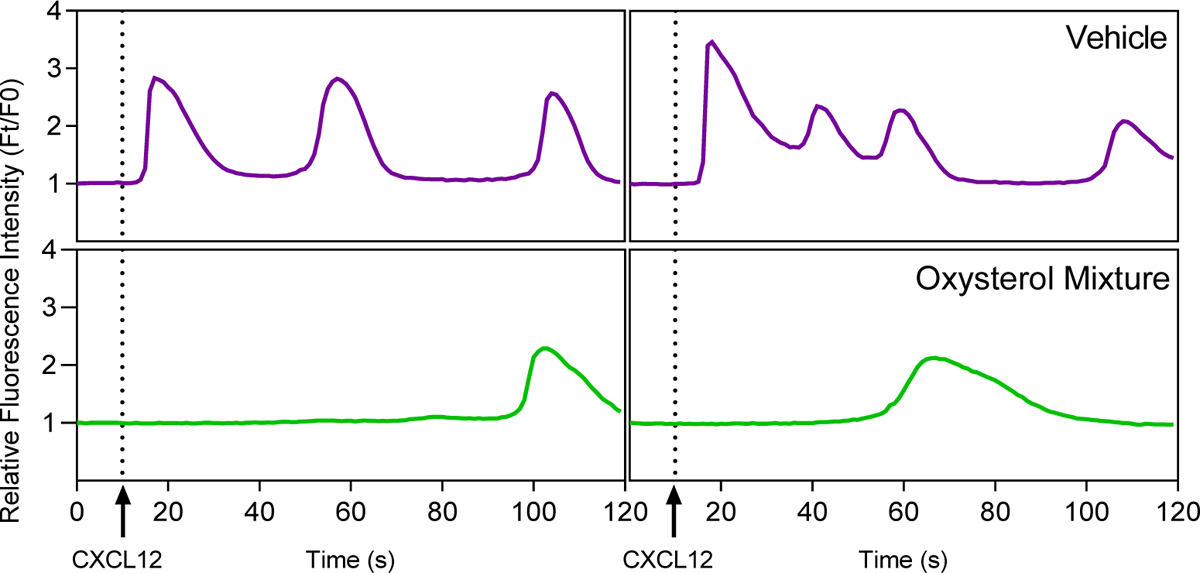
Calcium response in oxysterol treated cells. HeLa cells were treated with vehicle or oxysterol mixture for followed by calcium release analysis. Two single cell representatives of calcium response have been plotted.

**Figure S3:**
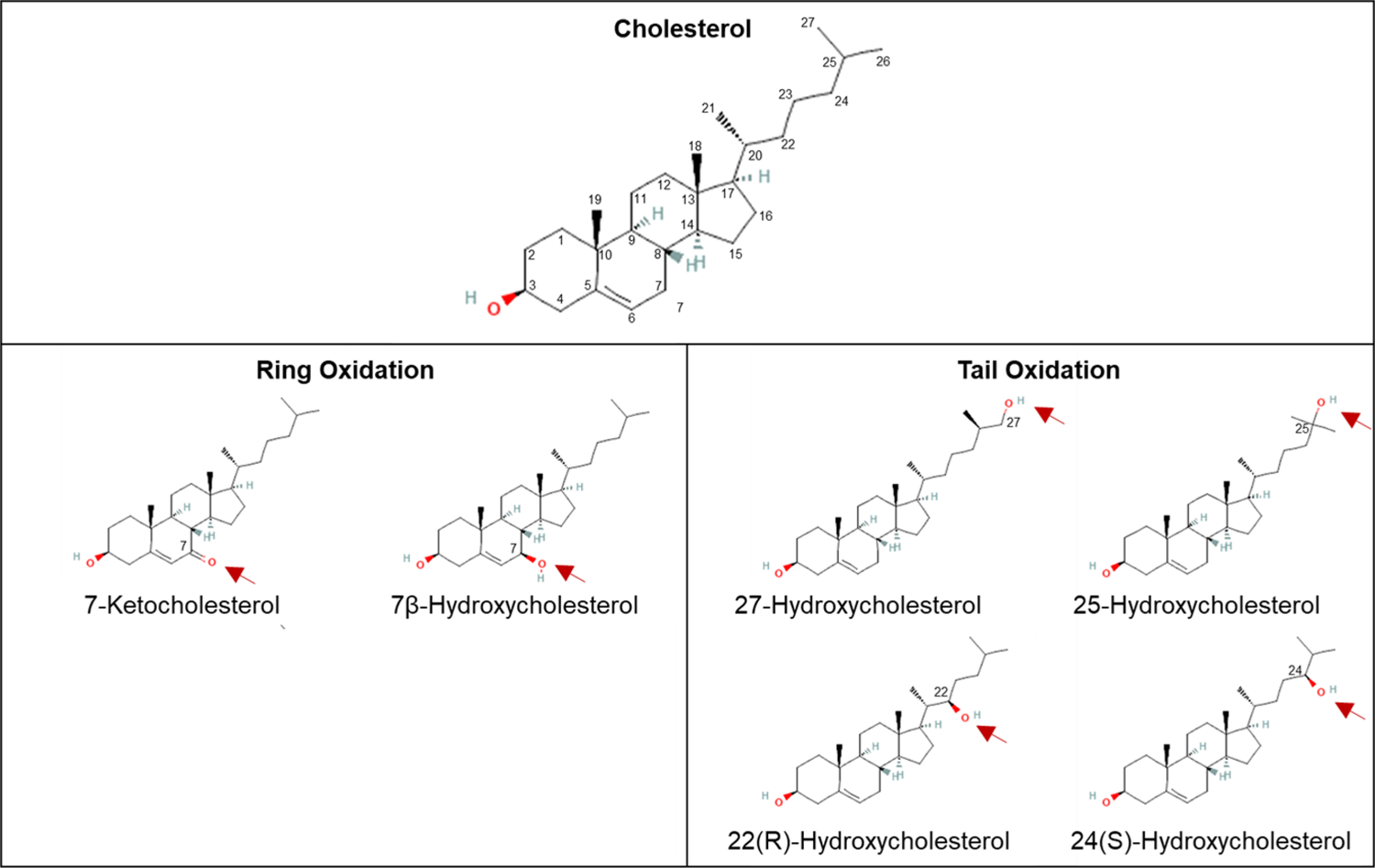
Structure of cholesterol and oxysterols used in this study. Structure of cholesterol (top) and its oxidized variants – oxysterols with the oxidation position marked in red. The oxysterols have been categorized into Ring (left panel) or Tail (right panel) oxysterols based on the position of oxidation.

**Figure S4:**
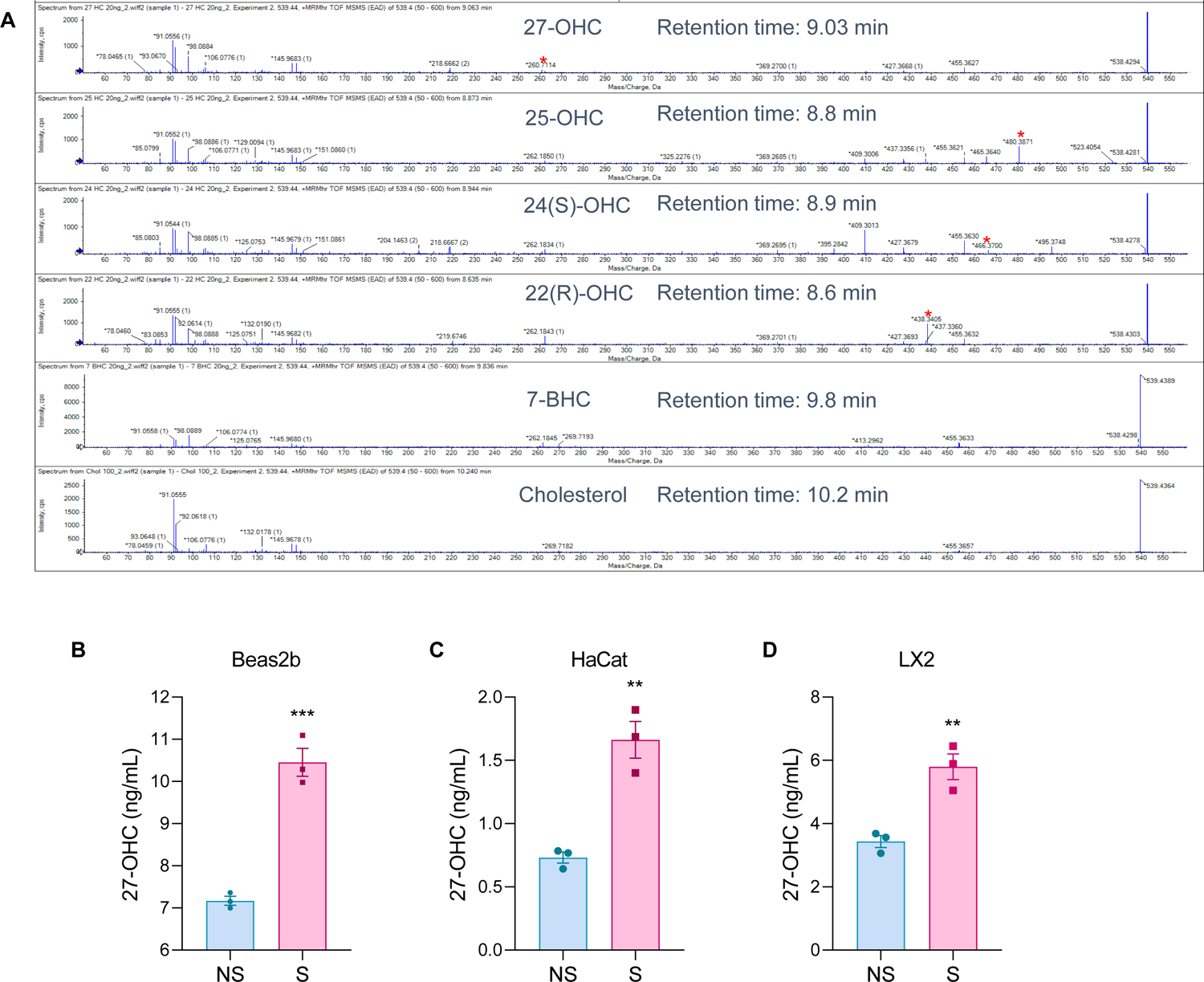
Oxysterol characterization and quantitation by LC-MS analysis. (a) Unique fragments used for analysis from cholesterol and oxysterol standards have been marked along with retention times. LC-MS analysis of 27-OHC in non-senescent and senescent primary cell lines of different tissue origin. Senescence was induced using ionizing radiation in all cells. (a) Beas2b, human bronchial epithelium cells (b) HaCat, human keratinocyte cells and (c) LX2, human hepatic stellate cells.

**Figure S5:**
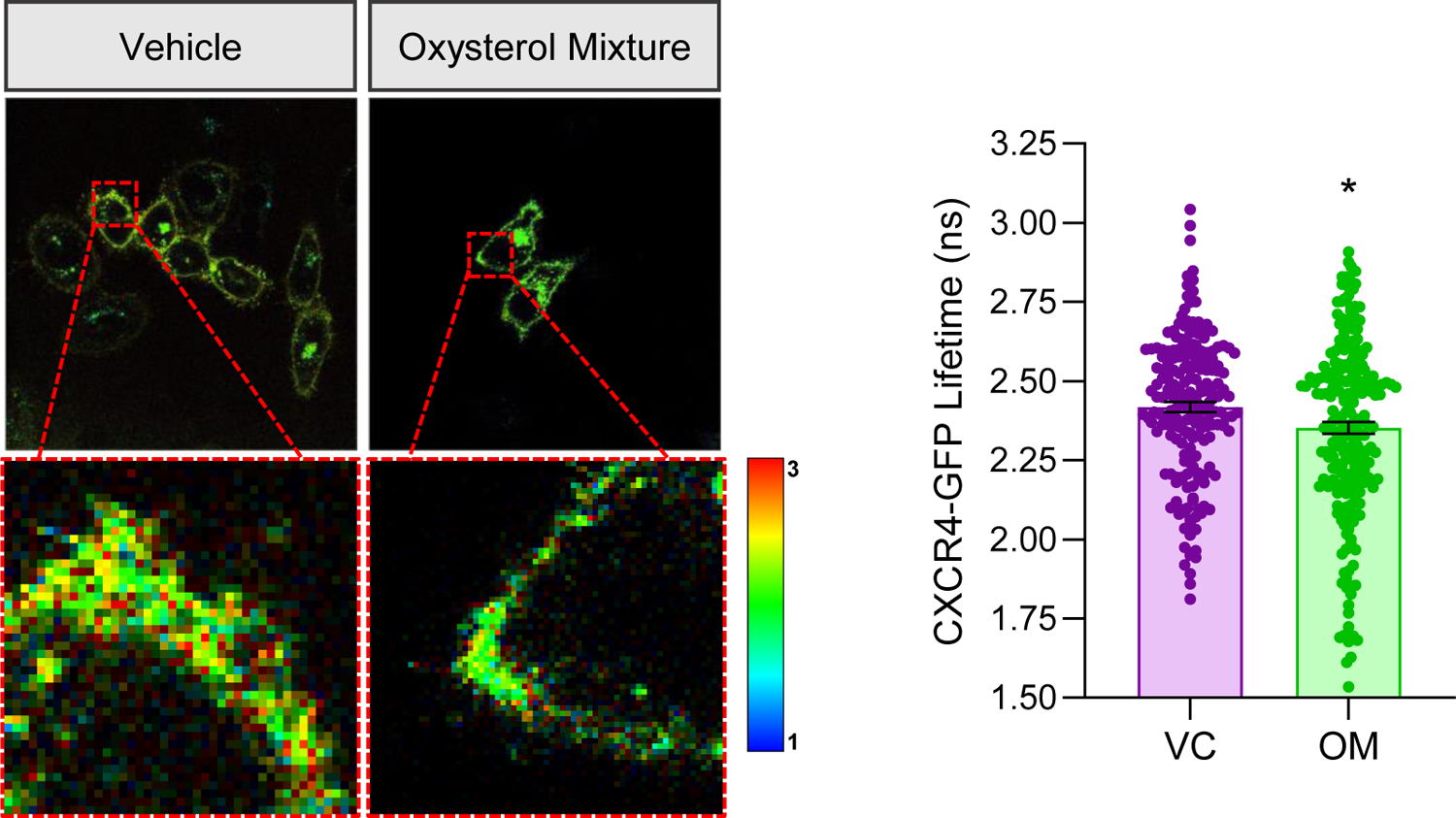
FLIM analysis for CXCR4 clustering. HeLa cells stably expressing CXCR4-GFP were treated with vehicle or oxysterol mixture followed by Fluorescence Lifetime Imaging (FLIM) to examine CXCR4 oligomerization. (A) Representative lifetime images for different groups and (B) Distribution of lifetime values.

**Figure S6:**
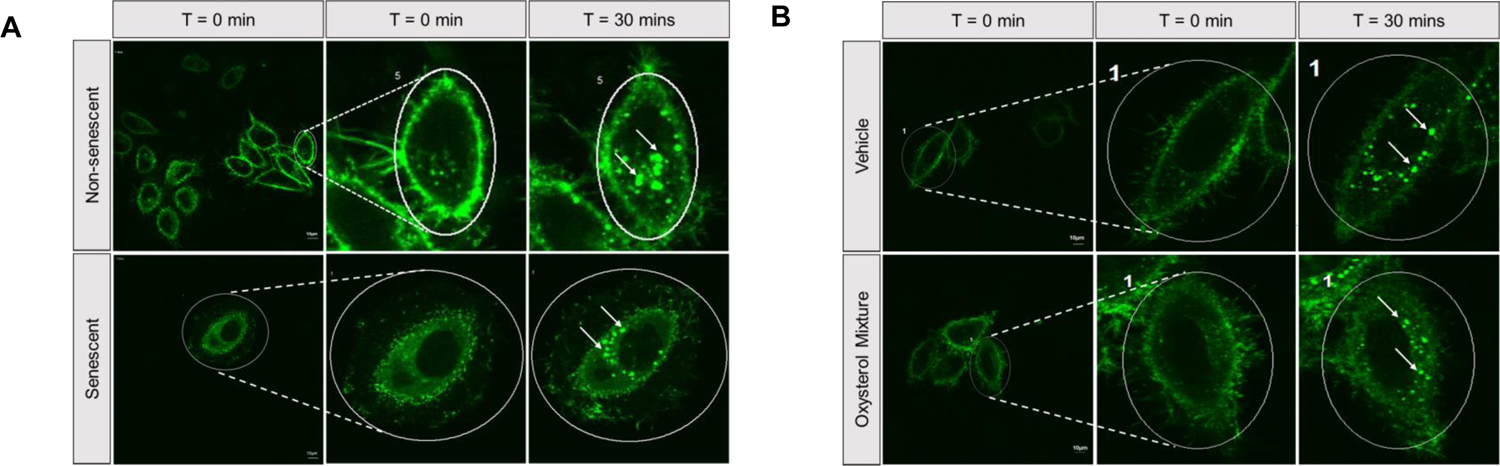
CXCR4 receptor internalization analysis after stimulation. HeLa cells stably expressing CXCR4-GFP were used for studying receptor internalization. (a) Non-senescent and senescent cells and (b) Non-senescent cells treated with vehicle or oxysterol mixture for 1 hour. Representative images at various time points after CXCR4 receptor activation with CXCL12 stimulation are shown (n=3; N>30 for all)

**Figure S7:**
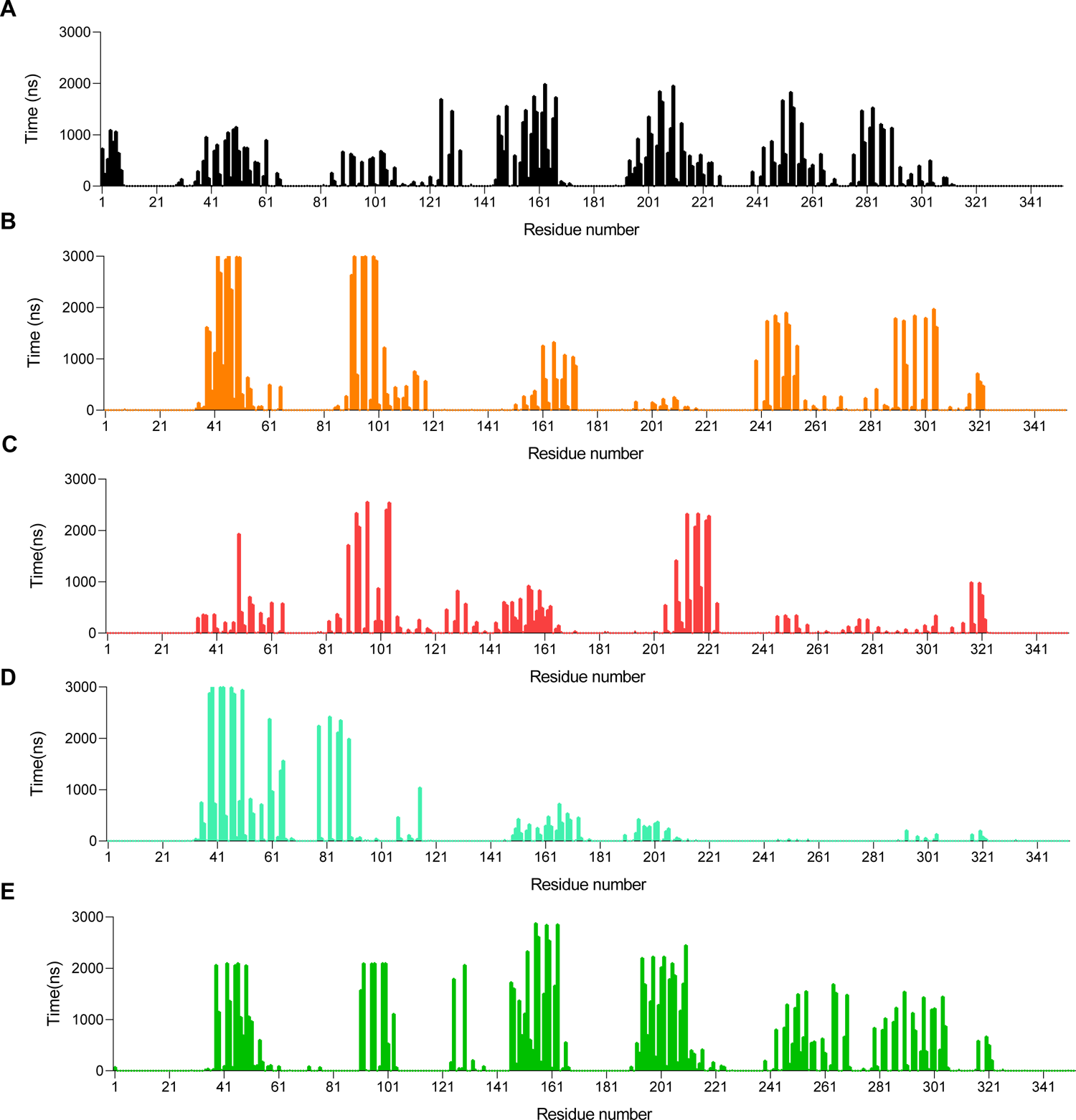
Binding time of various residues in CXCR4 receptor with sterols. Total binding time of the residues of CXCR4 with (a) cholesterol (b) cholesterol in CH-7BHC (c) 7-BHC in CH-7BHC (d) cholesterol in CH-27OHC and (e) 27-OHC in CH-27OHC, over 3μs.

**Figure S8:**
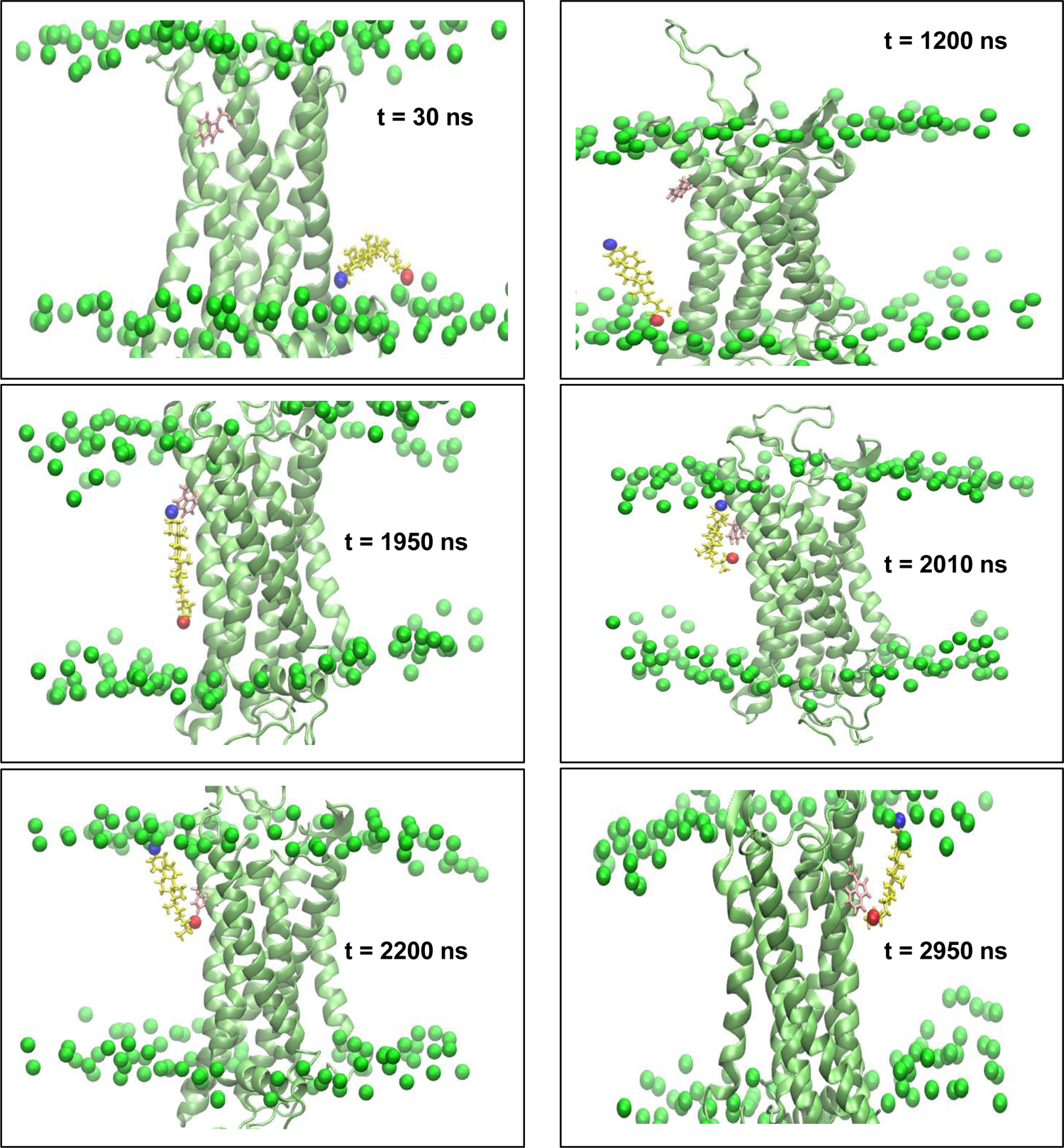
Various orientations of tail-oxidized sterol on membrane with CXCR4 receptor. Snapshots of a 27-OHC at different time points during the simulation illustrating different orientations and regions sampled. Here, CXCR4 is shown in lime, P atoms of the bilayer are shown as green spheres, residue W283 in pink and 27-OHC in yellow with its O3 as a blue sphere and O27 as a red sphere. At t = 30 ns, 27-OHC lies in a horizontal orientation, which is only observed with tail-oxidized sterols. At t = 1200 ns, it flips within the lower leaflet (a phenomenon never reported previously for any sterol), and at t = 1950 ns, it translocates to the upper leaflet driven by O3 interactions. At 2200 ns, O27 interacts with W283, and the molecule is now part of the upper leaflet, subsequently forming an H-bond with W283 at t = 2950 ns.

**Figure S9:**
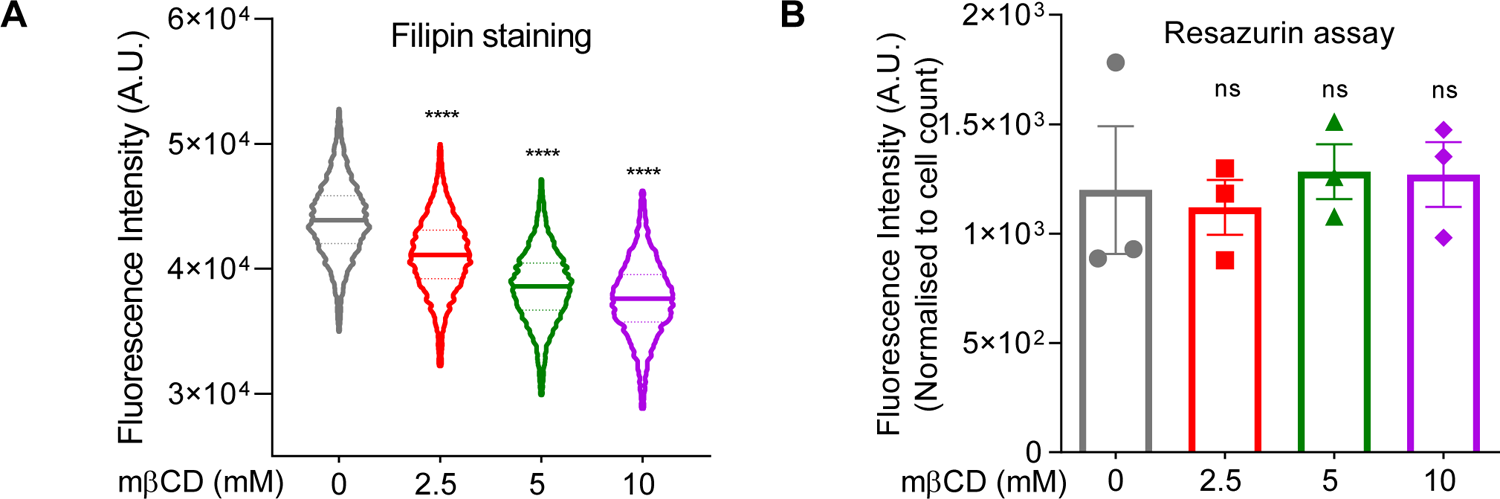
Cholesterol depletion assessment. Hela cells were treated with increasing concentrations of mβCD to deplete cholesterol. (a) Filipin staining to measure the cholesterol levels after mβCD treatment and (b) Resazurin assay to measure the metabolic activity after cholesterol depletion.

**Figure S10:**
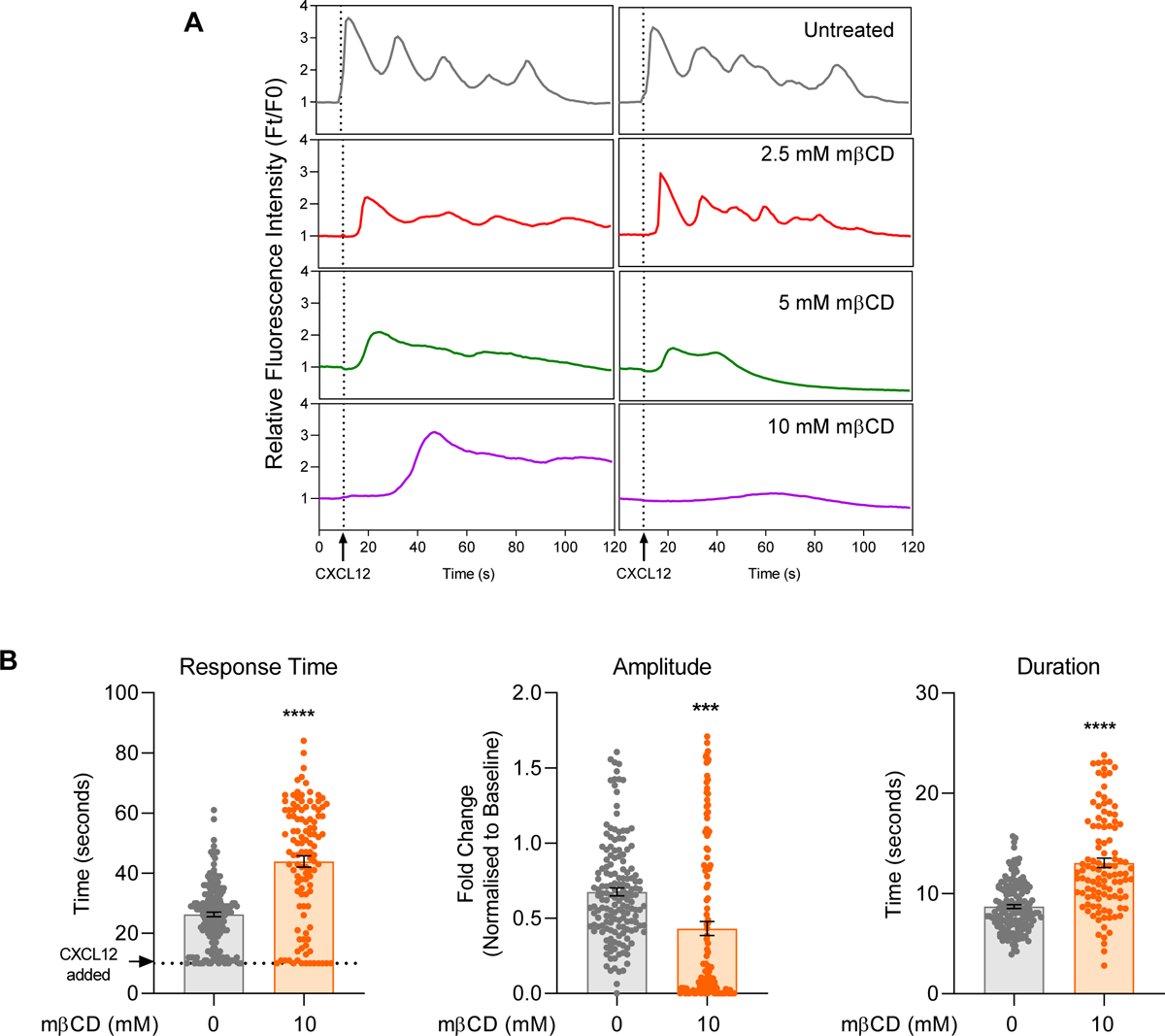
Single cell representations of calcium response after cholesterol depletion. Hela cells were treated with increasing concentrations of mβCD to deplete cholesterol followed by calcium release assay. (a) Two single cell calcium response representations are plotted over time for each treatment group. (b) Single cell analysis of untreated and 10mM mβCD treated cells for response time, amplitude and duration.

**Figure S11:**
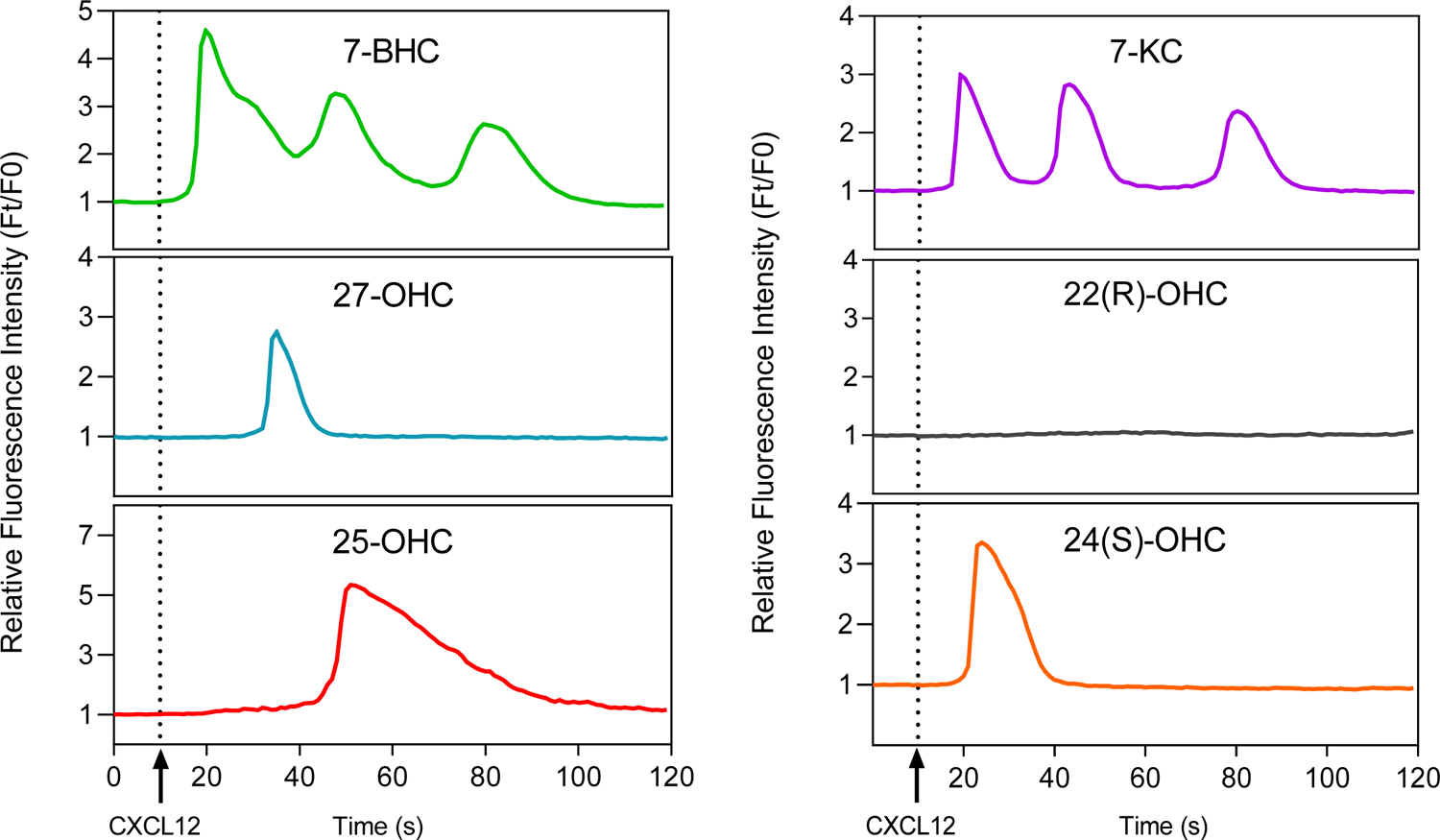
Single cell representatives of calcium response after oxysterol treatment. HeLa cells were treated with individual oxysterols (10µg/mL) for 1 hour followed by calcium release assay. A single cell representative for each oxysterol has been plotted with time.

**Figure S12:**
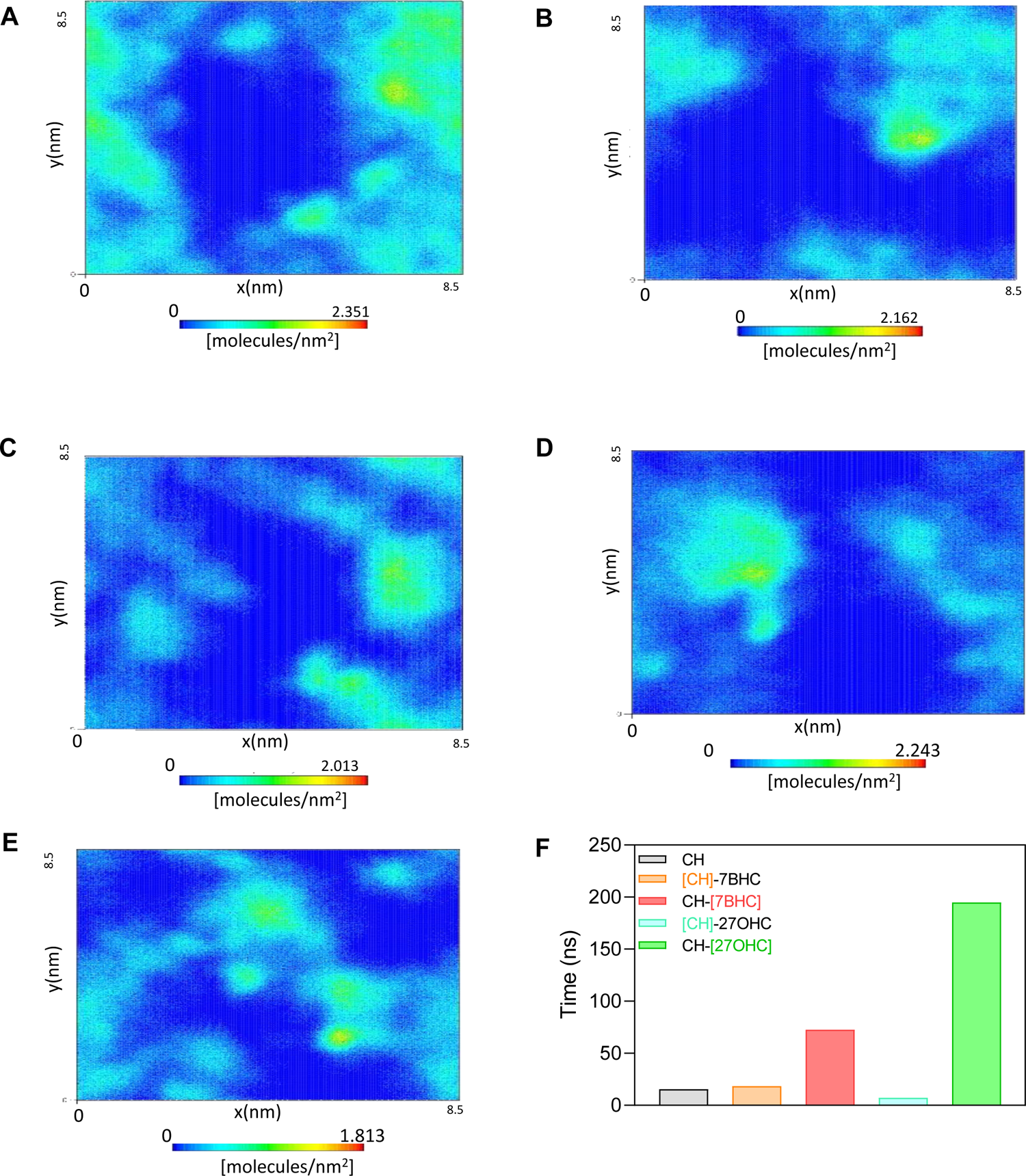
Demixing of 27-OHC and cholesterol. Two-dimensional number density map of (a) [CH], (b) [CH]-7BHC, (c) CH-[7BHC], (d) [CH]-27OHC and (e) CH-[27OHC], in the lower leaflet of the membrane for last 500 ns of the production run where x and y are the dimensions of the simulation box and the z-axis is normal to the membrane. Evidence for demixing tendency is observed in highly uncorrelated 2D density maps for cholesterol (d) and 27-OHC (e) in the CH-27OHC membranes. (f) Cumulative hydrogen bonding time of sterols with CXCR4 per sterol molecule. The distance cut-off for H-bonding was 3Å, and the angle cut-off used was 20°. Square brackets represent the specific sterol in the mixture.

**Figure S13:**
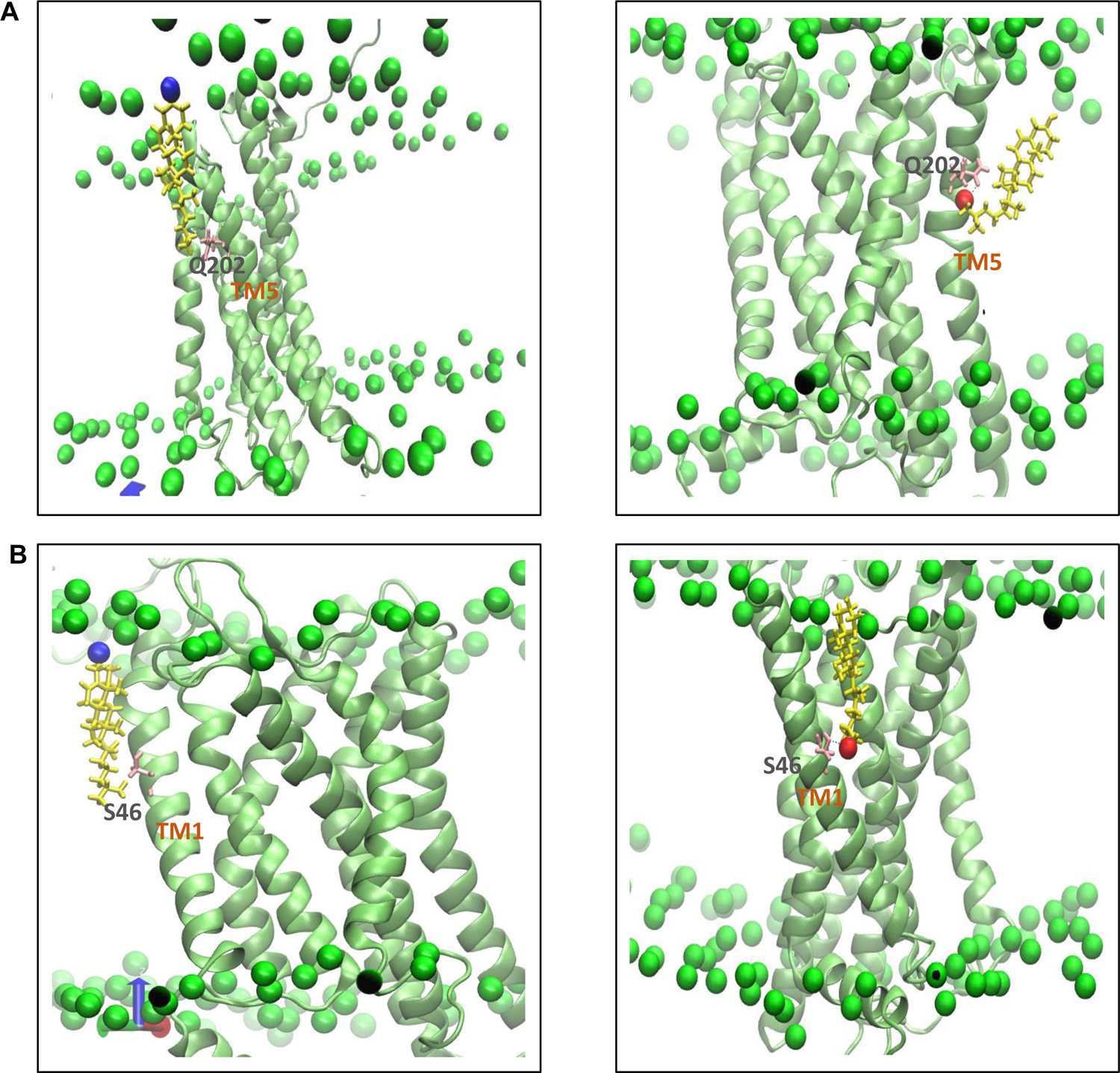
Presence of extra -OH group contributes to enhanced binding of tail-oxidized sterols to CXCR4. Snapshots highlighting that O27 present in 27-OHC drives the enhanced binding with 27-OHC compared to cholesterol. Here, CXCR4 is shown as in lime, P atoms of bilayer are shown as green spheres, residue concerned (Q202 and S46) in pink and sterol molecule in yellow with its O3 as blue sphere and O27 as red sphere. (a) cholesterol binds to Q202 with its hydrophobic tail in CH (left panel) and is replaced by 27-OHC in CH-27OHC (right panel) where O27 forms a H-bond with Q202, leading to longer and stronger binding (b) Cholesterol molecule in CH interacts with S46 in TM1 with its hydrophobic tail (left panel), which is replaced by 27-OHC in CH-27OHC (right panel) where O27 forms a very strong H-bond with S46 lasting for about 47% of the trajectory.

**Figure S14:**
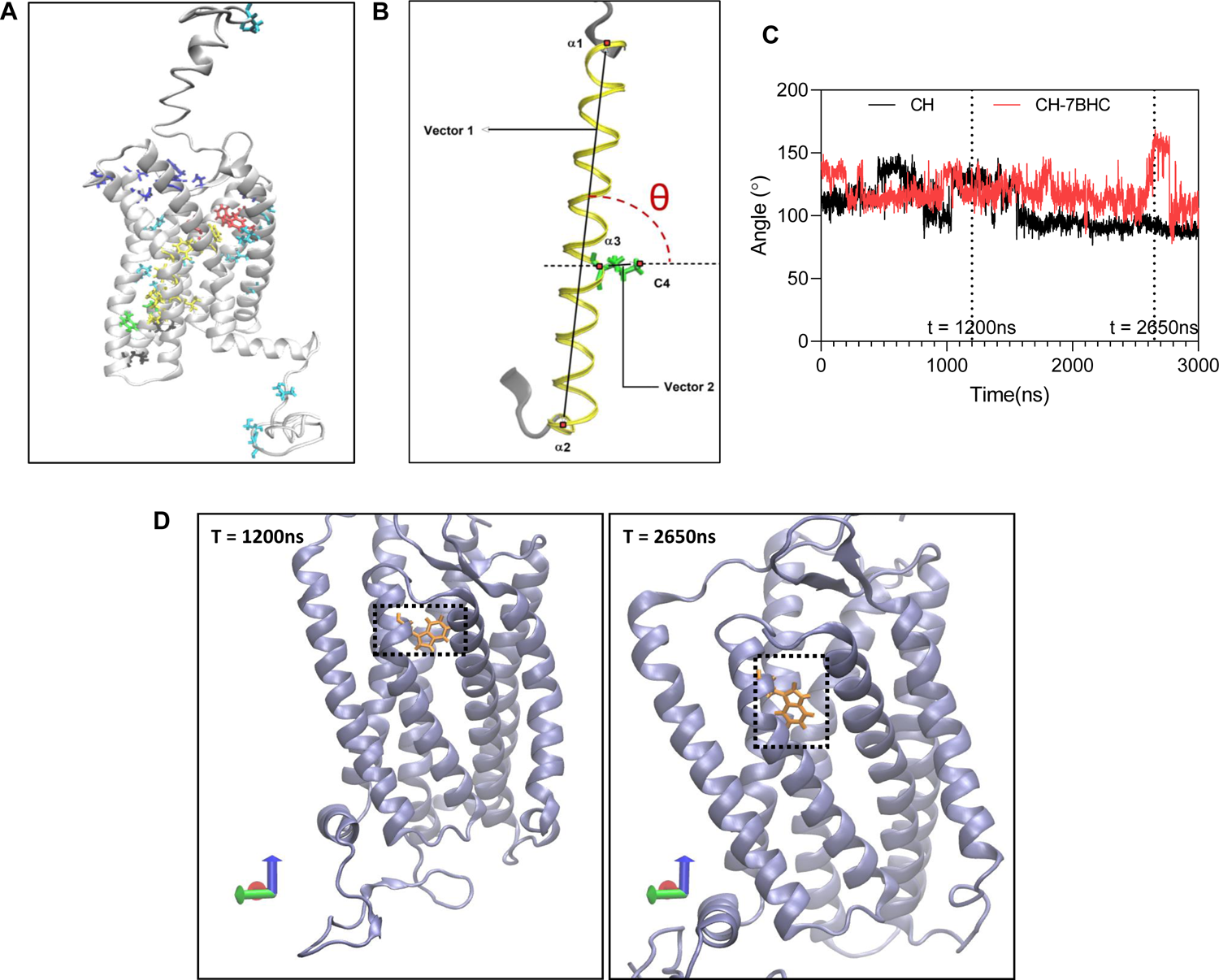
Orientation analysis of critical signalling residues in the presence oxysterols. (**a**) Critical signalling residues of the CXCR4 with chemokine engaging residues in blue, signal initiating residues in red, signal propagating residues in yellow, activation micro-switches in green, G-protein coupled residues in black and un-assigned residues in cyan (b) An illustration of the angle theta (θ), evaluated for critical signalling residues. (c) Time evolution of θ for W94 in CH and CH-7BHC systems, (d) Illustration of two orientations of W94 in CH-7BHC corresponding to different values of θ at different time points in the simulations.

**Figure S15:**
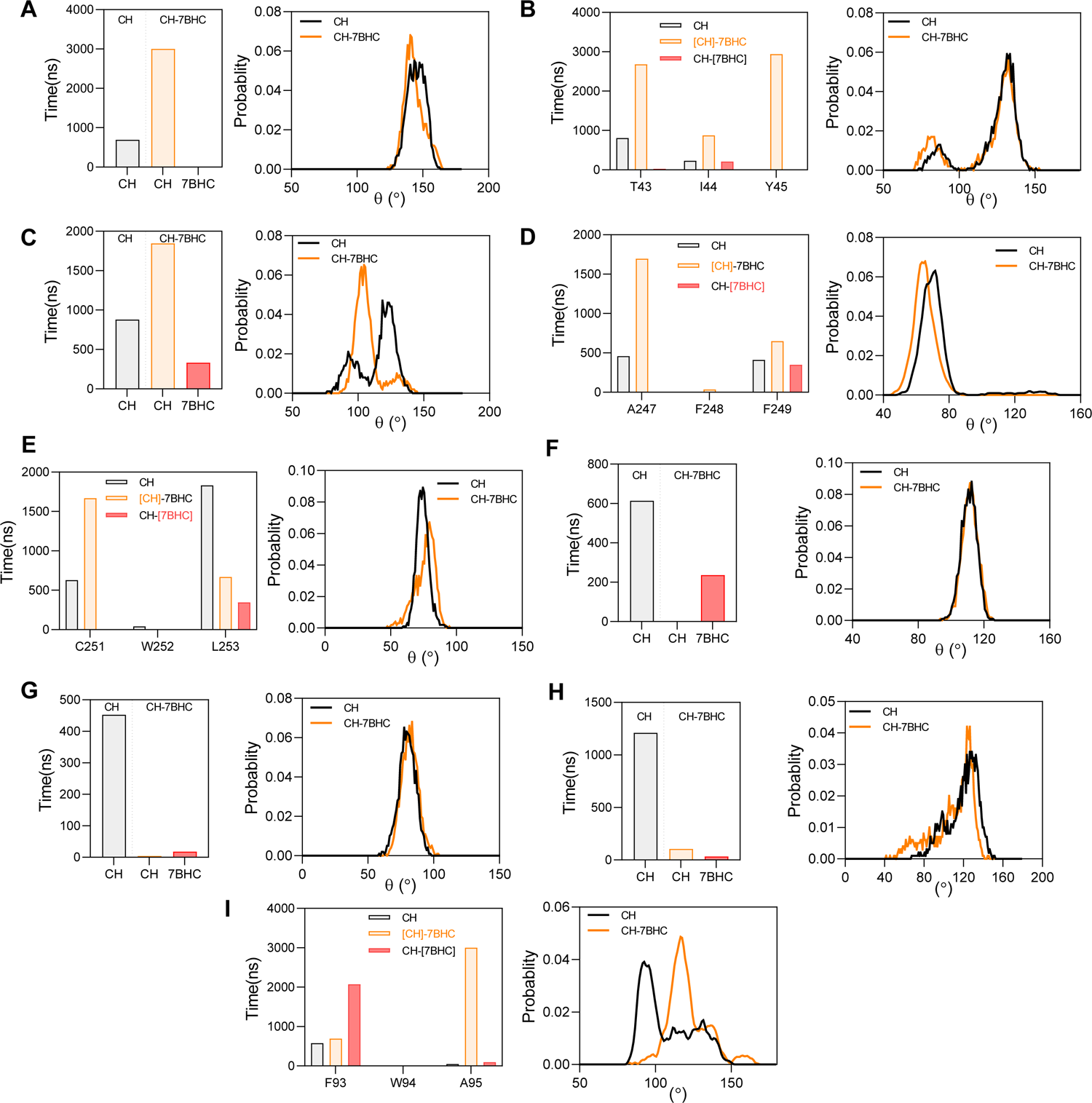
Changes in orientations of critical residues in presence of 7-BHC. (a-i) Effect of sterol interactions on critical signalling residues, where change in their interaction time compared to CH is shown in the left panel and change in θ is illustrated in the right panel. C1 interactions are illustrated in (a) P42, (b) I44, (c)L246, (d) F248 and (e) W252, and C2 interaction in (f) A128, (g) Y219 and (h) I286, and C4 interaction in (i) W94, in CH-7BHC. See main text for criterion used to define C1-C4 interactions.

**Figure S16:**
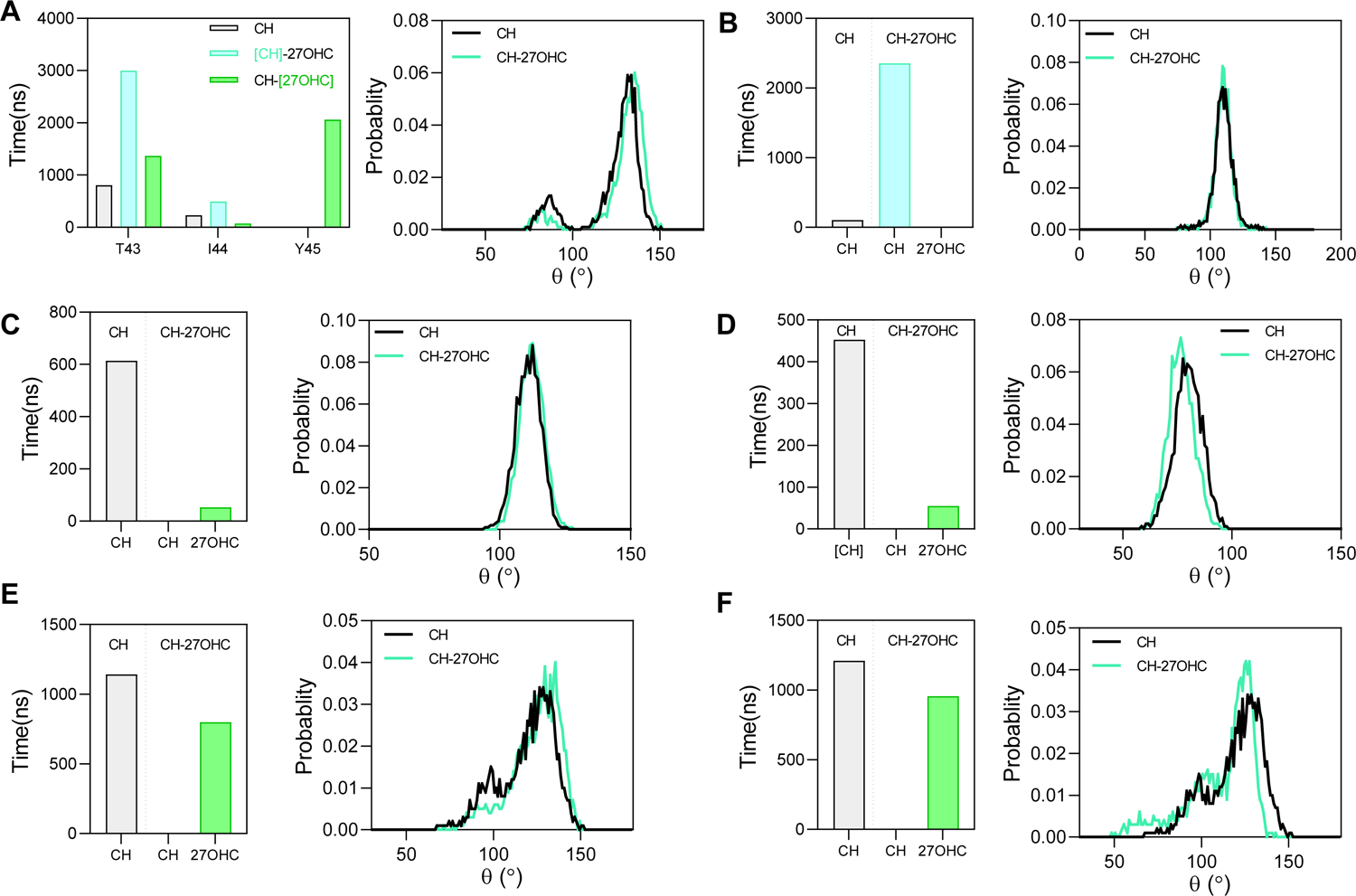
Changes in orientations of critical residues in the presence of 27-OHC. Effect of sterol interactions on critical signalling residues where change in their interaction time compared to CH is shown in the left panel and change in θ is illustrated in the right panel. C1 interactions for (a) I44 and (b) L86, C2 interaction for (c) A128 and (d) Y219, and C3 interactions as (e) K282 and (f) I286, in CH-27OHC. See main text for criterion used to define C1-C4 interactions.

**Figure S17:**
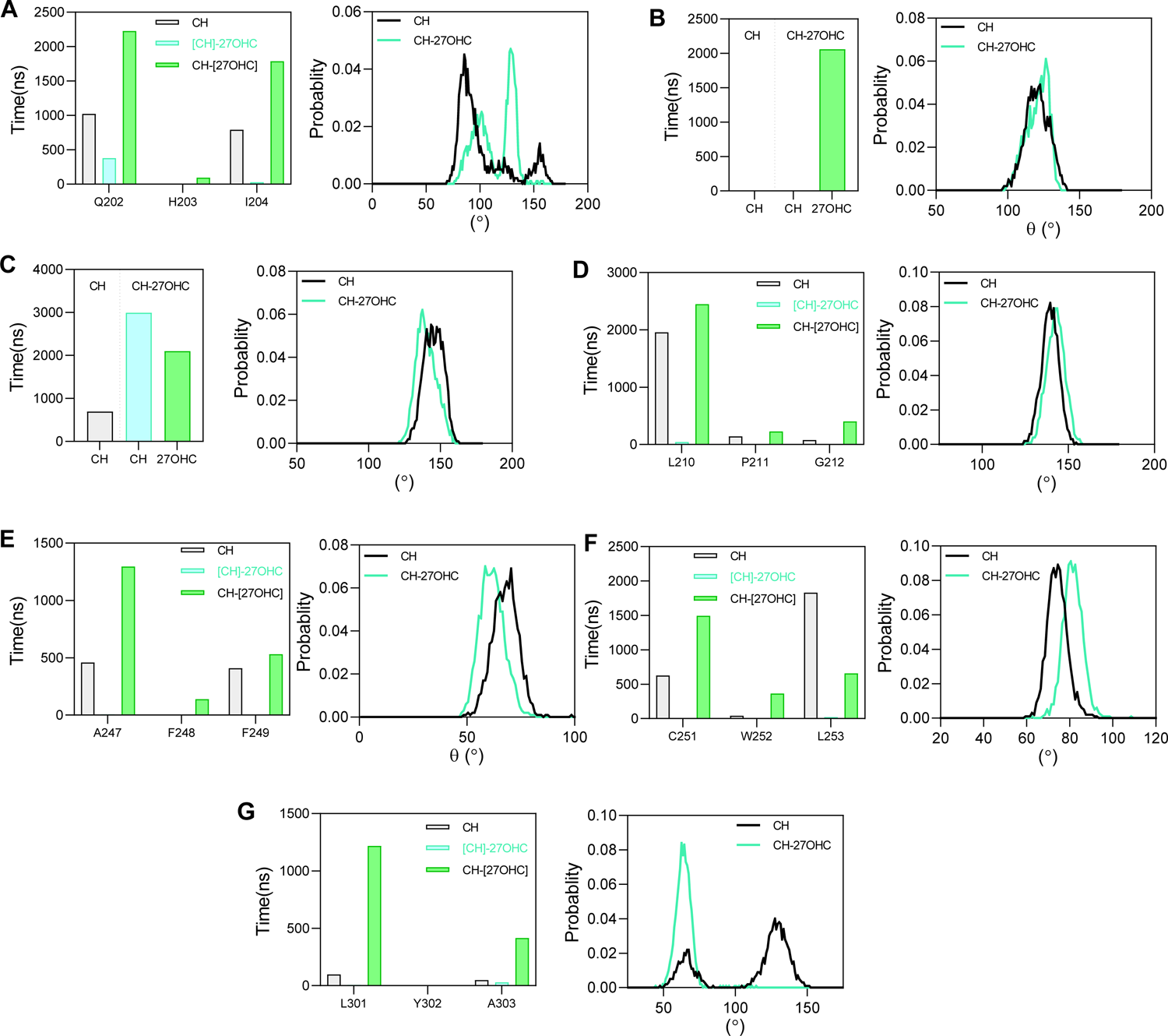
Changes in orientations of critical residues due to enhanced binding with 27-OHC. Effect of sterol interactions on critical signalling residues where a change in their interaction time compared to CH is shown in the left panel and change in θ is illustrated in the right panel. C4 interactions as (a) H203, (b) Y45, (c) P42, (d) P211, (e) F248, (f) W252, and (g)Y302, in CH-27OHC. See main text for criterion used to define C1-C4 interactions.

**Figure S18:**
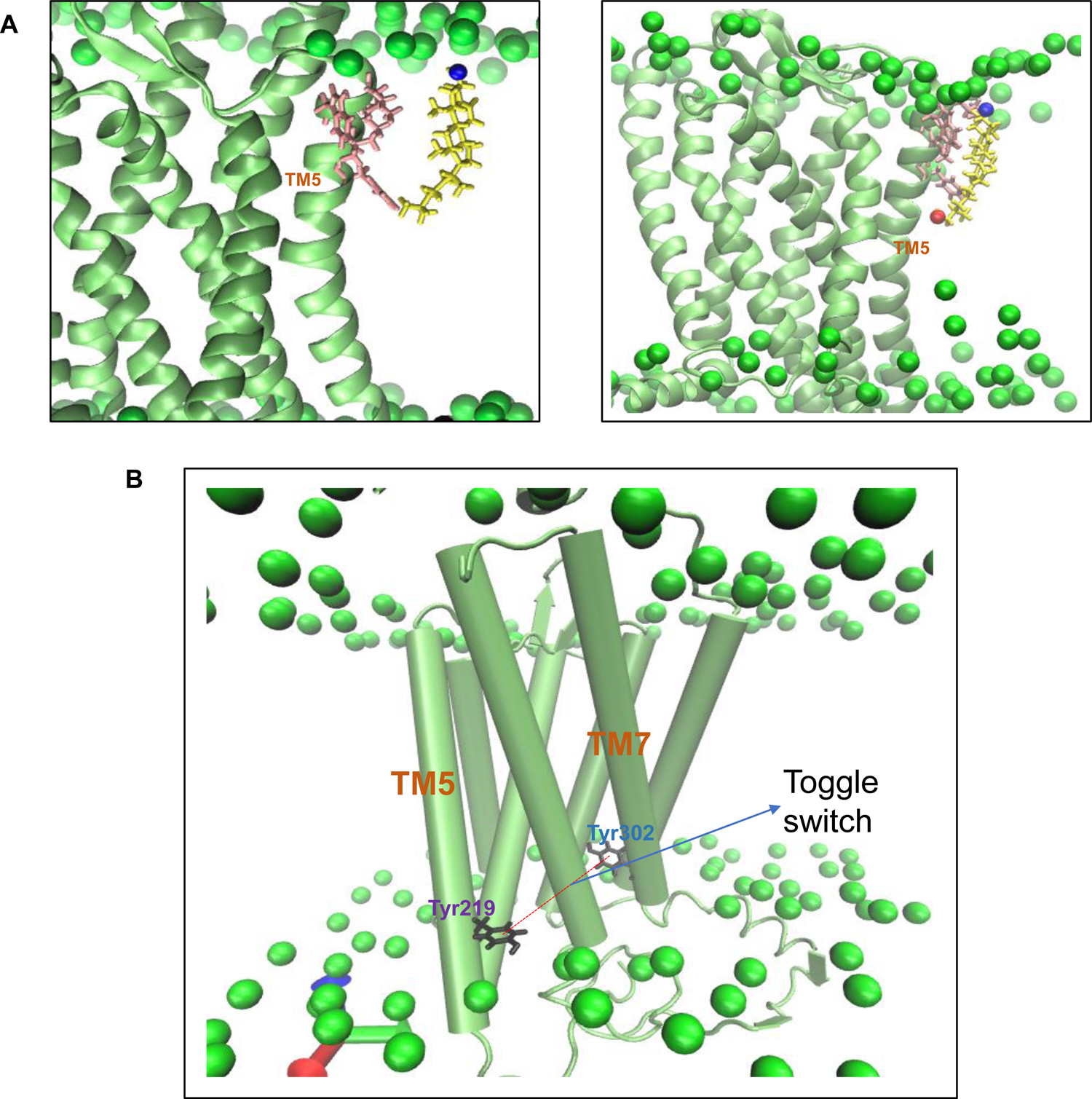
Snapshots illustrating (a) dimeric interface and (b) toggle switch in CXCR4 protein. CXCR4 is shown in lime, P atoms of the bilayer are shown as green spheres, residues of the dimeric interface in pink and sterol molecule in yellow with its O3 as blue sphere and O27 as red sphere. (a) Cholesterol molecule interacts with the dimeric interface (L194, W195, V197, V198 and F201) in CH (left panel). It is replaced by 27-OHC in CH-27OHC (right panel). 27-OHC binds for a much longer time than cholesterol. (b) Toggle switch which is the distance between Y219 and Y302 is illustrated with residues Y219 and Y302 shown in black.

**Figure S19:**
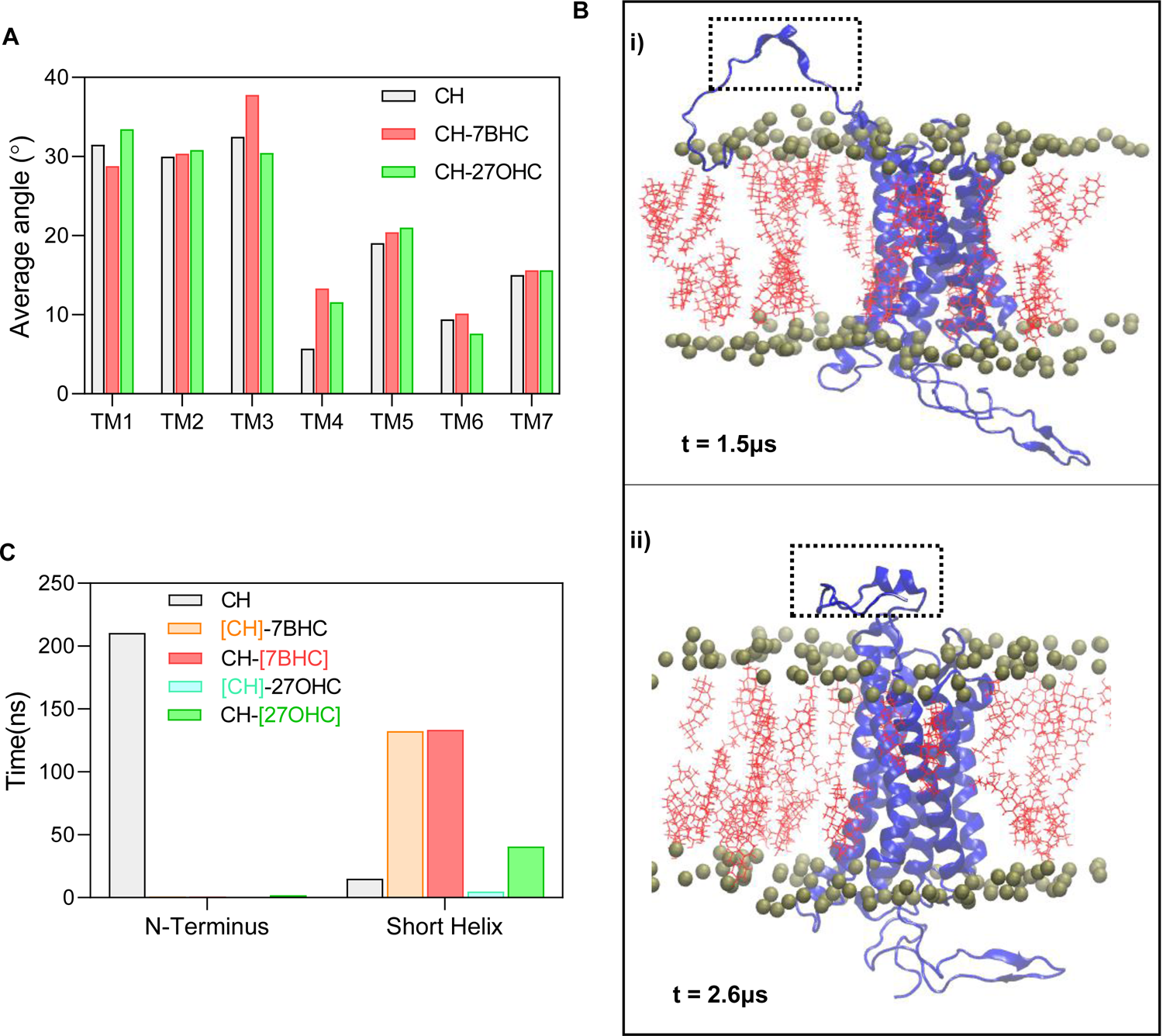
Tilt angle of TMs and changes at N and C terminus in presence of oxysterols. (a) Tilt angle of all the TMs of CXCR4 calculated as the angle between the vector passing through the first and last C-α atom of a TM and a vector normal to the membrane, averaged over 3μs (b) snapshots of N-terminus of CXCR4 in CH system at t = 1500 ns (b-i) where it enters the membrane and interacts with the cholesterol molecules and at t = 2600ns (b-ii) where it forms a helix and is present in the extracellular region. Here, CXCR4 is shown in blue, cholesterol molecules in red and P atoms of the bilayer are shown in metal (c) Cumulative hydrogen bonding time of sterols with initial ten residues of N-terminus of CXCR4 which interact with the membrane in all three systems and in the starting twenty residues of C-terminus (short helix) which form the short eighth-helix, calculated as the cumulative time spent by the residues in hydrogen bonding with sterol molecules.

**Figure S20:**
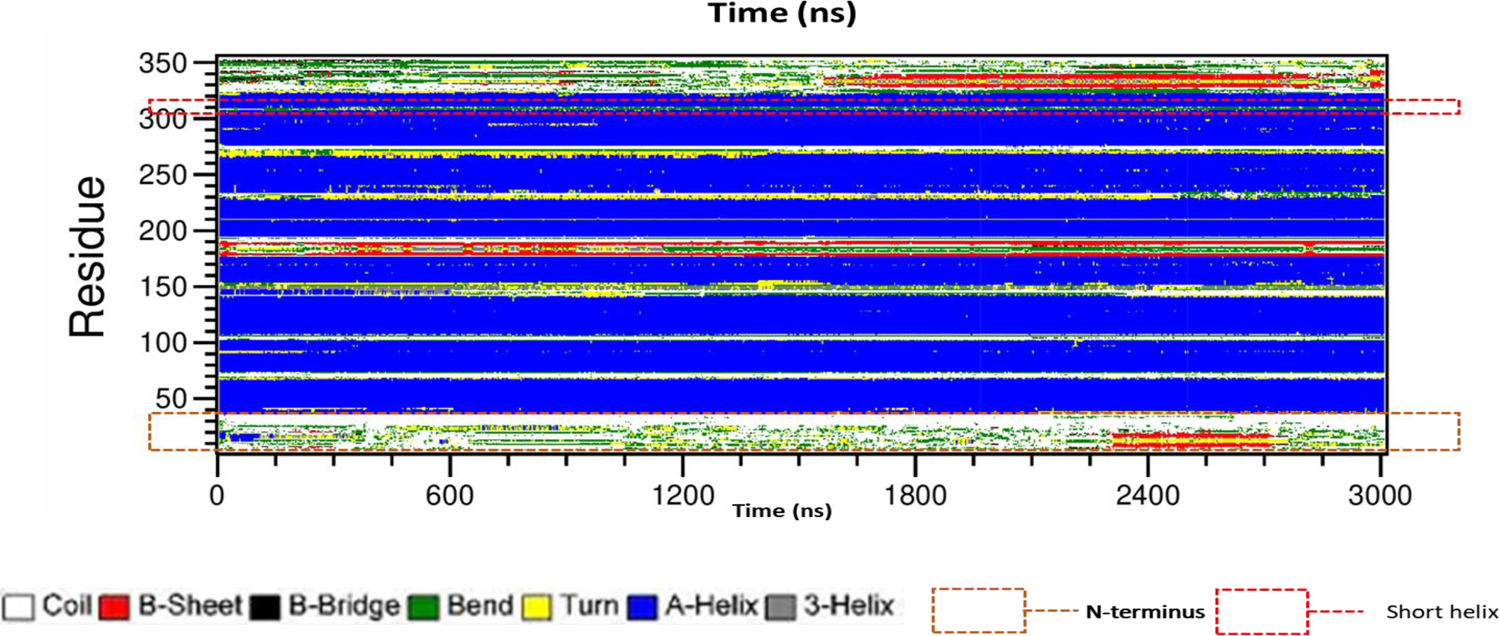
Conformational changes in CXCR4. Secondary structure prediction of CXCR4 with time over 3 μs in CH-27OHC.

**Figure S21:**
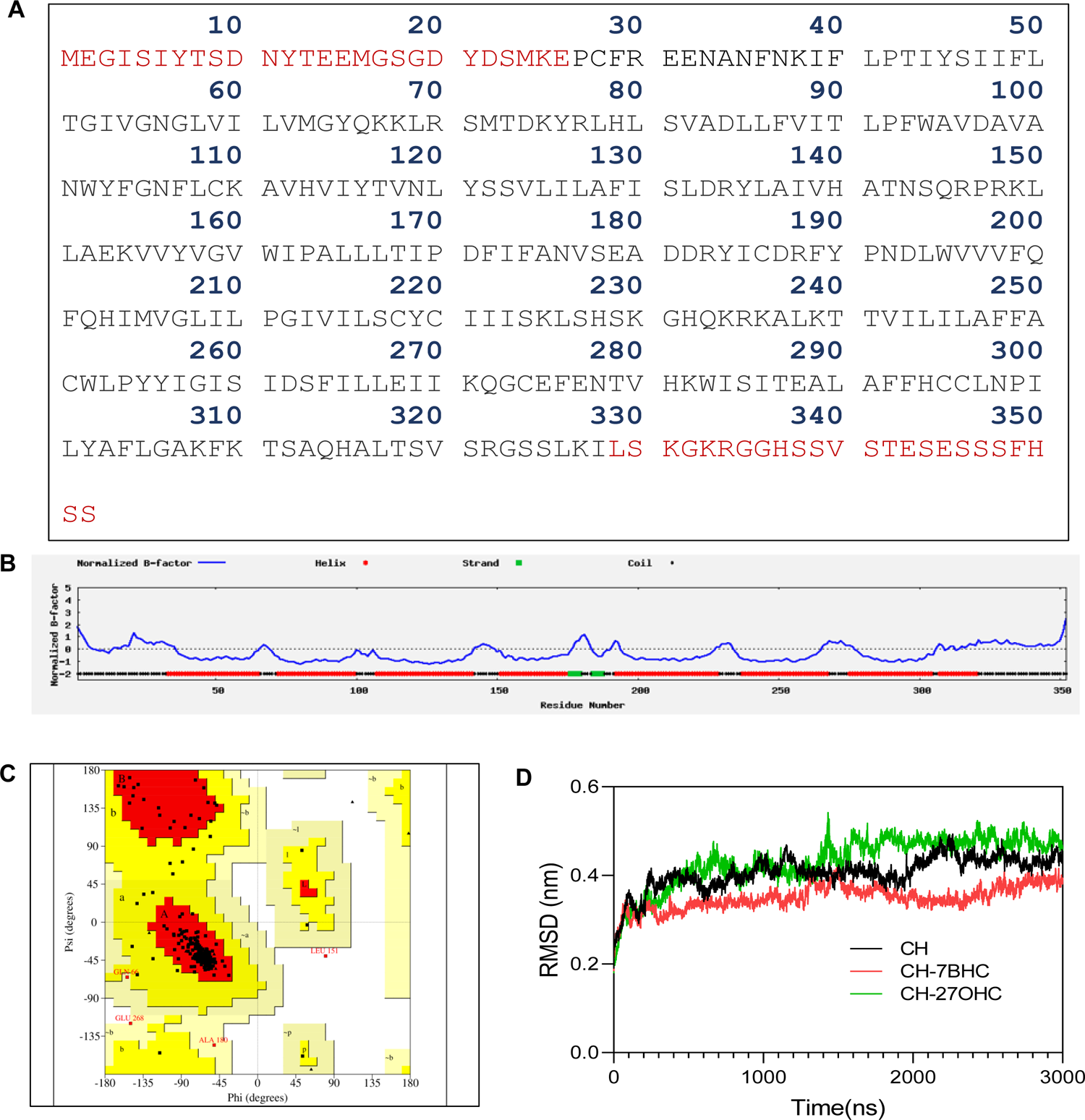
Analysis of stability of the model structure. (a) Sequence of amino acids of CXCR4 with residues missing in PDB entry 3ODU highlighted in red (b) The normalized B-factor of the residues of the model structure (c) Ramachandran plot of residues of CXCR4 inside the membrane in CH system, at the end of the simulation. Region A, B and L are the most favoured, a, b, l and p are the additionally allowed regions, ∼a, ∼b, ∼l and ∼p are the generously allowed regions, and the rest is the disallowed region (d) root mean square displacement (RMSD) of CXCR4 calculated from its initial position for the 3μs simulation in all three systems.

**Figure S22:**
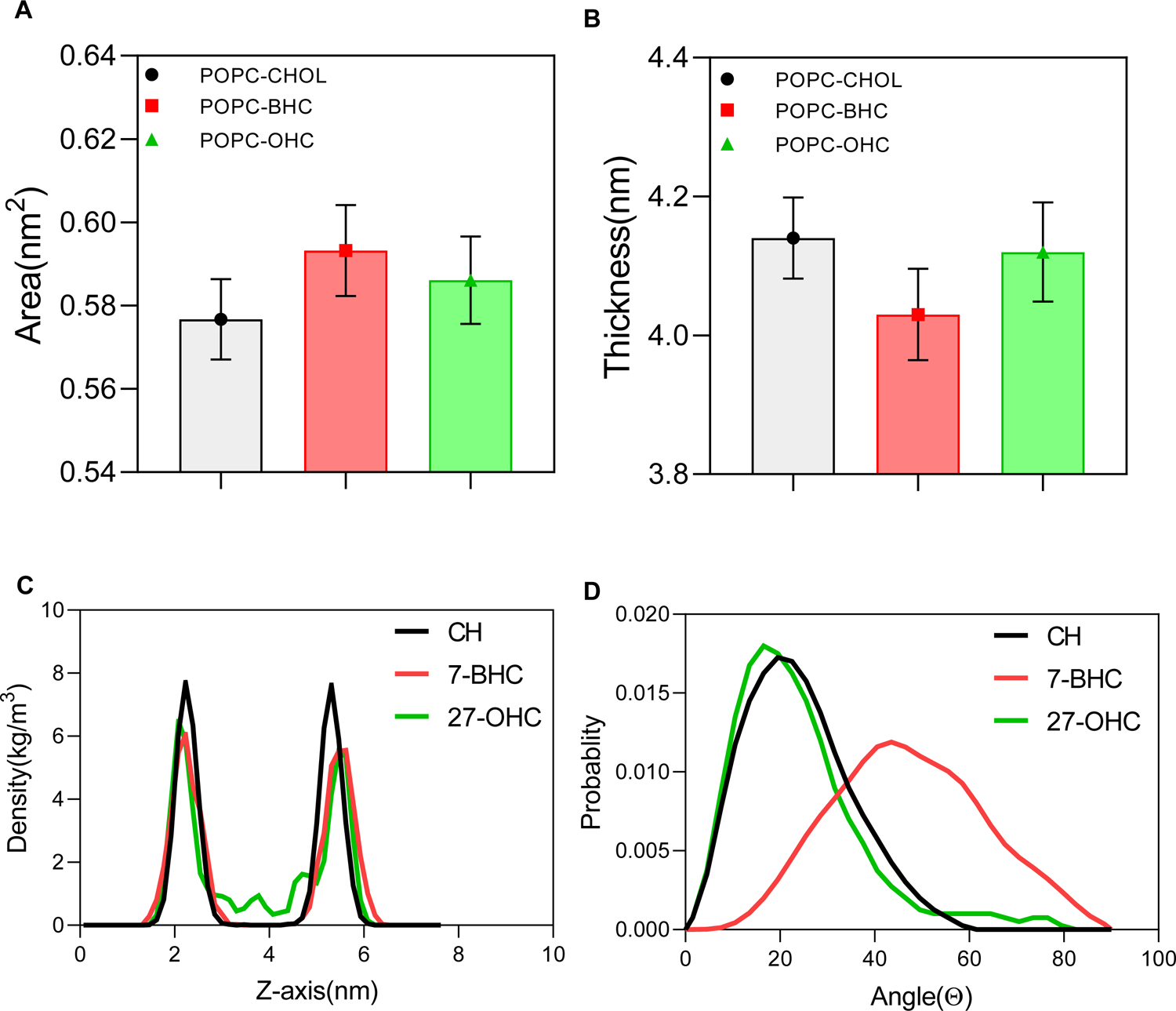
Bare membrane simulations. (a) Area per molecule of the membrane calculated as the area of the simulation box divided by the number of molecules (POPC + sterols) in each leaflet. (b) Thickness of the membrane averaged obtained by calculating the distance between phosphorous atoms of the upper and lower leaflet. (c) Average number density of the oxygen atom O3 along the axis normal to the membrane. (d) Tilt angle distribution of the sterol molecules calculated as the angle between the vector passing through C3 and C27 of the sterol molecules and the vector normal to the membrane.

---

**Movie S1. Calcium oscillations in non-senescent cells after stimulation with CXCL12.** Calcium release analysis in non-senescent HeLa cells depicting oscillations in single cells post-stimulation with CXCL12. The ligand was added at the 10^th^ frame capture, and calcium response was recorded.

**Movie S2. Calcium oscillations in senescent cells after stimulation with CXCL12.** Calcium release analysis in BrdU treated senescent HeLa cells depicting oscillations in single cells post stimulation with CXCL12. The ligand was added at the 10^th^ frame capture, and calcium response was recorded.

**Movie S3. Calcium oscillations in non-senescent cells after vehicle treatment and stimulation with CXCL12.** Calcium release analysis in non-senescent HeLa cells treated with vehicle (ethanol) depicting oscillations in single cells post-stimulation with CXCL12. The ligand was added at the 10^th^ frame capture, and calcium response was recorded.

**Movie S4. Calcium oscillations in non-senescent cells after depleting cholesterol and stimulation with CXCL12.** Calcium release analysis in non-senescent HeLa cells treated with 10 mM mβCD depicting oscillations in single cells post-stimulation with CXCL12. The ligand was added at the 10^th^ frame capture, and calcium response was recorded.

**Movie S5. Calcium oscillations in non-senescent cells after treatment with oxysterol mixture and stimulation with CXCL12.** Calcium release analysis in HeLa cells treated with oxysterol mixture depicting oscillations in single cells post stimulation with CXCL12. The ligand was added at the 10^th^ frame capture, and calcium response was recorded.

**Movie S6. Calcium oscillations in senescent cells after treatment with NAC and stimulation with CXCL12.** Calcium release analysis in senescent HeLa cells treated with 10mM NAC for 24 hours followed by stimulation with CXCL12. The ligand was added at the 10^th^ frame capture, and calcium response was recorded

**Movie S7. Calcium oscillations in senescent cells after treatment with Quercetin and stimulation with CXCL12.** Calcium release analysis in senescent HeLa cells treated with 2µM Quercetin for 24 hours followed by stimulation with CXCL12. The ligand was added at the 10^th^ frame capture, and calcium response was recorded.

